# Novel exported bifunctional fusion enzymes with chorismate mutase and cyclohexadienyl dehydratase activity: shikimate pathway enzymes teamed up in no man’s land

**DOI:** 10.1101/2023.03.13.532365

**Authors:** Christian Stocker, Tamjidmaa Khatanbaatar, Kathrin Würth-Roderer, Gabriele Cordara, Ute Krengel, Peter Kast

## Abstract

Chorismate mutase (CM) and cyclohexadienyl dehydratase (CDT) catalyze two subsequent reactions in the intracellular biosynthesis of phenylalanine. Surprisingly, exported CMs and CDTs exist in bacterial pathogens. Here, we report the discovery of novel and extremely rare exported bifunctional fusion enzymes, consisting of fused CM and CDT domains. Such enzymes were found in only nine bacterial species belonging to non-pathogenic γ- or β-proteobacteria. In γ-proteobacterial fusion enzymes, the CM domain is N-terminal to the CDT domain, whereas in β-proteobacteria the order is inversed. The CM domains share 15-20% sequence identity with the AroQ_γ_ class CM holotype of *Mycobacterium tuberculosis* (*MtCM), and the CDT domains 40-60% identity with the exported monofunctional enzyme of *Pseudomonas aeruginosa* (PheC). *In vitro* kinetics revealed a *K*_m_ <7 µM, much lower than for *MtCM, whereas kinetic parameters are similar for CDT domains and PheC. There is no feedback inhibition of CM or CDT by the pathway’s end product Phe, and no catalytic benefit of the domain fusion compared to engineered single-domain constructs. The fusion enzymes of *Aequoribacter fuscus*, *Janthinobacterium* sp. HH01, and *Duganella sacchari* were crystallized and their structures refined to 1.6, 1.7, and 2.4 Å resolution, respectively. Neither the crystal structures nor size-exclusion chromatography show evidence for substrate channeling or higher oligomeric structure that could account for cooperation of CM and CDT active sites. The genetic neighborhood with genes encoding transporter and substrate binding proteins suggests that these exported bifunctional fusion enzymes may participate in signaling systems rather than in the biosynthesis of Phe.

## Introduction

Chorismate mutase (CM) catalyzes the rearrangement of chorismate to prephenate, which in turn is converted to phenylpyruvate by prephenate dehydratase (PDT), a cyclohexadienyl dehydratase (CDT) specific for prephenate (Figure 1). Both CM and CDT are crucial enzymes for the biosynthesis of phenylalanine (Phe) in plants, fungi, and bacteria *via* the shikimate pathway.^1, 2^ The activities of key enzymes in this pathway are tightly controlled because its products are energetically expensive, and the reactions are closely linked with other metabolic processes.^1^ Control over the metabolic flux in this pathway is typically regulated in bacteria by feedback inhibition of the first enzyme of the pathway, 3-deoxy-D-*arabino*-heptulosonate 7-phosphate (DAHP) synthase,^3^ and at the branch point of the pathway, by regulating the activity of CM, often in conjunction with an associated PDT or a fused prephenate dehydrogenase (PDH) domain, and of anthranilate synthase.^4–6^

**Figure 1.**
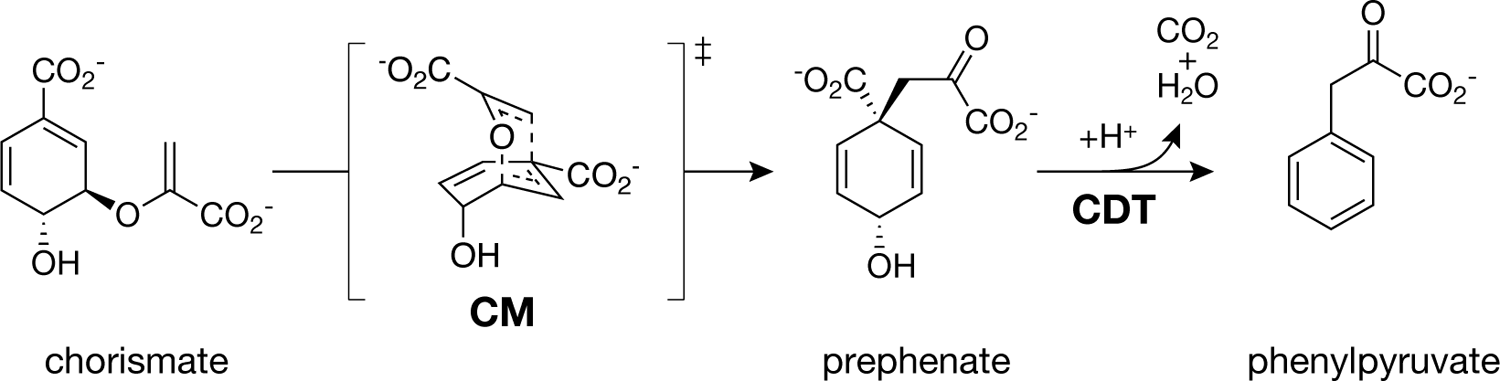
Reactions catalyzed by chorismate mutase and cyclohexadienyl dehydratase. Chorismate mutase (CM) catalyzes the Claisen rearrangement of chorismate to prephenate *via* an oxabicyclic transition state. Cyclohexadienyl dehydratase (CDT) thereafter converts prephenate to phenylpyruvate.

Two families of CMs, the AroH and AroQ class, are known, which exhibit evolutionarily and structurally distinct folds. The predominant family found in nature is the AroQ class, which can be further divided into 4 subclasses AroQ_α_ to AroQ_δ_. AroQ CMs have α-helical folds,^1, 7–11^ with the AroQ prototype, the AroQ_α_ CM from *Escherichia coli* (EcCM), consisting of two intertwined protomers forming a 6-helix bundle with two active sites. To enable metabolic pathway regulation AroQ CMs frequently occur as bifunctional fusion enzymes or are involved in protein complexes. CMs fused to a PDT domain usually belong to the AroQ_α_ subclass and are found in the cytoplasm. For example, PheA in *E. coli* (also called P-protein) consists of the AroQ_α_ CM domain EcCM fused to both a PDT domain and a third regulatory domain, forming a fusion protein found in most proteobacteria.^9, 12, 13^ The CM and PDT activities of PheA are reduced by 55% and 85%, respectively, in the presence of excess L-Phe, the relevant end product of the biosynthetic pathway.^14, 15^ Furthermore, *E. coli* also entails an AroQ_α_ CM domain fused to a PDH, the bifunctional fusion enzyme TyrA (also called T-protein) that is also common in enteric bacteria.^12, 13^ Unlike PheA, TyrA does not have a third regulatory domain; instead, feedback regulation in TyrA occurs through L-tyrosine (Tyr) binding near the active site of PDH, leading to full inhibition of PDH activity, whereas CM retains 93% of its activity.^5^

Alternative modes of metabolic feedback-control are known for the intracellular CMs from the AroQ_ý_ and AroQ_δ_ subclasses. The CM of *Saccharomyces cerevisiae* is of the AroQ_ý_ subclass and forms a dimer composed of 12 α-helices.^8^ Each subunit harbors a catalytic center, which is activated by Trp and inhibited by Tyr upon binding to different allosteric sites at the interface of the two protomers.^16^ AroQ_δ_ CMs only occur in the cytoplasm of the taxonomic class *Actinobacteria* and include the well-studied enzymes of *Mycobacterium tuberculosis* and *Corynebacterium glutamicum*.^17^ The dimeric AroQ_δ_ CMs strongly resemble AroQ_α_ consisting of two protomers of three α-helices each. Whereas AroQ_δ_ by itself shows <1% of the activity of a typical CM, it reaches its full potential upon forming a complex with its partner enzyme, the DAHP synthase.^10^ Complex formation allows for inter-enzyme allosteric control of CM activity *via* binding sites for the three aromatic amino acids on the DS.^17^ In *M. tuberculosis*, the binding of Phe and Tyr inhibit intracellular CM activity by 72% and 37%, respectively.^10^ In the CM-DS enzyme system from *C. glutamicum,* metabolic feedback regulation by the aromatic amino acids is even more pronounced.^18^

AroQ_γ_ subclass CMs are very peculiar in that they are exported from the cytoplasm, the compartment where the shikimate pathway is localized. Exported CMs (*CMs) are mainly found in bacteria, but also present in fungi and nematodes.^19–21^ The best characterized AroQ_γ_ family member is *MtCM from *M. tuberculosis*. It forms a homodimer, with each protomer consisting of six α-helices. Whereas the four N-terminal helices contribute the active site, a disulfide bond in the C-terminal part between helices 5 and 6 is thought to be essential for structural integrity when exported out of the cell.^9^ Notably, no feedback regulation was detected for *MtCM.^19, 22^ Since biosynthesis of intracellular metabolites outside of the cell is not plausible, AroQ_γ_ enzymes were hypothesized to be candidates for virulence factors, especially for mammalian pathogens, as they are also found in *Salmonella enterica* serovar Typhimurium*, Pseudomonas aeruginosa*, and *Yersinia pestis*.^13, 19, 22–24^ This hypothesis is supported by the finding that *CMs from several phytopathogenic bacteria, fungi, and nematodes are involved in host invasion.^20, 25–29^

*P. aeruginosa* also produces an exported CDT enzyme, PheC, which in contrast to its cytoplasmic PDT counterpart,^30, 31^ is not feedback regulated.^32^ PheC consists of two α/β subdomains connected by two flexible hinge strands, with the ligand binding site located at the interface of the two subdomains. Based on structural homology, PheC probably evolved from periplasmic solute-binding proteins and was shown to oligomerize to homodimers or homotrimers.^30, 32^ Since *P. aeruginosa* also produces an exported CM and maybe even an exported aromatic aminotransferase, which together could form a complete metabolic path from chorismate to phenylalanine,^13^ it has been speculated that an excess of internally produced chorismate escaping the cell is captured and converted by this so-called ‘hidden overflow pathway’.^32, 33^ However, the biological role of these exported enzymes is still controversially discussed.

Here we report the discovery of novel and highly rare exported bifunctional fusion enzymes with *CM and CDT domains, that appear to exist solely in non-pathogenic bacteria and should thus have another function than virulence. Extensive kinetic, biophysical, and structural characterization and genetic neighborhood analysis are employed to gain insights in their functioning and shed some light on their biological role.

## Results

### Highly rare, exported fusion enzymes with CM and CDT domains

The full amino acid sequence of the AroQ_γ_ CM from *M. tuberculosis* (*MtCM),^19^ including its N-terminal secretion signal (UniProt: P9WIB9), was taken as input for a BLASTP search in the NCBI database.^34^ Besides other monofunctional exported CMs with >80% sequence coverage, a few search hits stood out with 50% (or less) sequence coverage due to the presence of an additional protein domain. A further BLASTP search of this other domain revealed that it shared >40% identity with the exported cyclohexadienyl dehydratase PheC of *P. aeruginosa* (*PaCDT). Furthermore, two distinct topologies of these exported bifunctional fusion enzymes were found; either the CM domain is N-terminally fused to a C-terminal cyclohexadienyl dehydratase domain (*CMCDT), or *vice-versa* the CDT domain is N-terminal and the CM domain C-terminal (*CDTCM). The novel exported fusion proteins appear to be extremely rare and are only found in very few organisms, most of which belonging to the taxonomic group of γ-proteobacteria. These mostly aquatic species encompass *Shewanella baltica* (Sb), the main culprit for producing hydrogen sulfide on rotting fish,^35–37^ and *Shewanella psychrophila* (Sp),^38^ both *Thalassomonas actiniarum* (Ta) and *Thalassomonas viridans* (Tv), which were suggested to be part of the natural microflora of sea anemone,^39^ *Aequoribacter fuscus* (Af), which belongs to the family *Halieaceae* that accounts for more than 10% of the total ocean surface bacterioplankton,^40, 41^ and *Steroidobacter cummioxidans* (Sc), a rubber degrader that secretes oxygenases to cleave polyisoprenes (Figure 2).^42^

**Figure 2.**
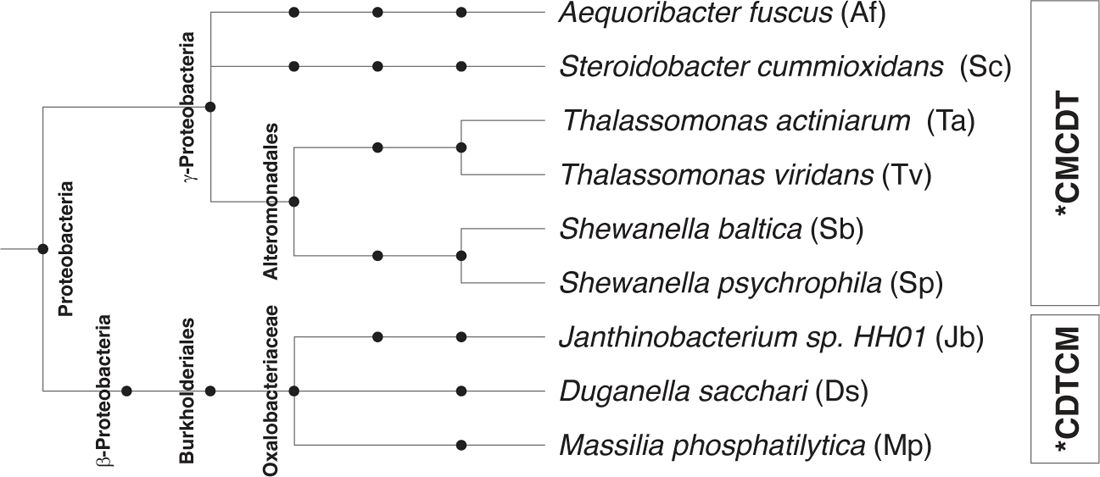
Phylogenetic tree representation of the exported *CMCDT and *CDTCM fusion proteins. Shown is the phylogenetic relationship between the proteobacterial species that possess an exported *CMCDT or *CDTCM bifunctional fusion enzyme. Each node represents a taxonomic sub-classification.

*CDTCM sequences are even rarer than *CMCDTs, and solely found in very few species of the class of β-proteobacteria. These include *Duganella sacchari* (Ds), isolated from the rhizosphere of sugar cane,^43^ *Massilia phosphatilytica* (Mp), a genus closely related to the genus *Duganella* based on 16S rRNA sequence comparisons^43^ and capable of solubilizing phosphate from rocky fertilizers,^44^ and the *Janthinobacterium* sp. HH01 (Jb) from a biofilm-forming family known to synthesize antibacterial and antifungal compounds.^45^

The clear division between exported *CMCDTs occurring only in γ-proteobacteria and exported *CDTCMs only found in β-proteobacteria (Figure 2) speaks against horizontal gene transfer between the two taxonomic groups with respect to these exported bifunctional fusion enzymes. More likely, these genes evolved from different and independent primordial protein fusion events, by convergent evolution.

### Comparison of primary structures

The amino acid sequences of *MtCM and *PaCDT (PheC) were aligned with those of the exported *CMCDT (Figure 3) and *CDTCM (Figure 4) fusion enzymes to identify domain boundaries and conserved patterns, and to gain initial structural insights. Calculated sequence similarity and identity scores of the individual CM or CDT domains suggest folds similar to those of *MtCM and *PaCDT (Figure S1). Notably, generally, lower scores were obtained for the CM domains towards *MtCM (≈30-40% similarity, ≈15-25% identity) than for the CDT domains towards *PaCDT (≈55-75% similarity, ≈40-60% identity).

**Figure 3.**
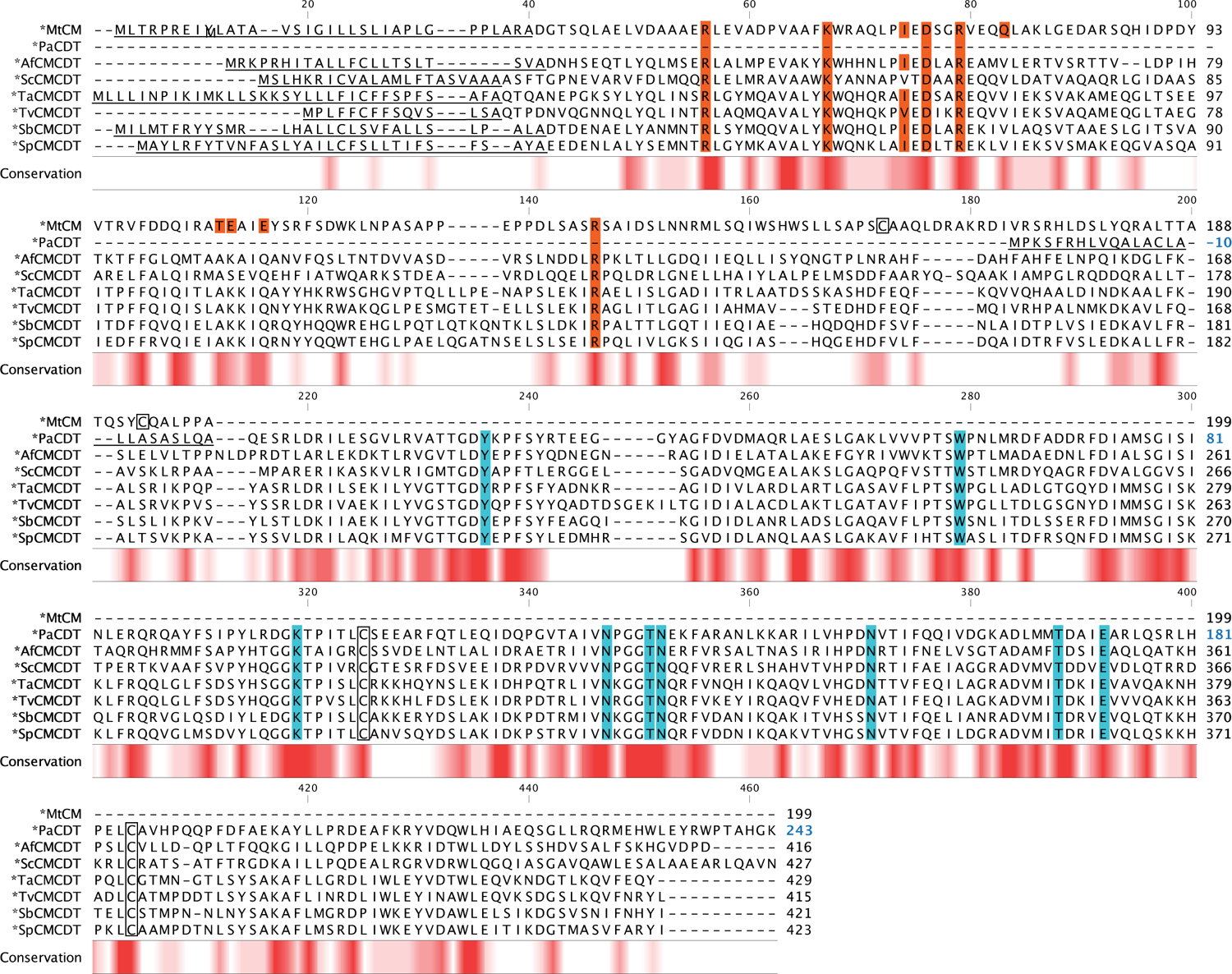
Sequence alignment of bifunctional *CMCDT variants. The amino acid sequences of the *CMCDT bifunctional fusion enzymes are aligned to *MtCM (AroQγ subclass) and *PaCDT (also called PheC). Signal sequences are underlined, and established *MtCM and *PaCDT active site residues (and their fully conserved counterparts in the new sequences) highlighted in red and blue, respectively. Cys residues forming a disulfide bond in *MtCM and *PaCDT are boxed. Residue numbering shown on the right starts with the first residue (Met1) of the open reading frame including the signal sequence, except for *PaCDT, where the initial residue corresponds to the N-terminus of the mature protein after signal sequence cleavage (Gln1). This exception, distinguished with blue numbers, allows for compatibility with the nomenclature adopted by Clifton *et al.* in their work on *PaCDT and thereby for consistency in the discussion of active site residues;^32^ it is also applied in the structural figures. The color gradients from white to dark red below the alignment indicate the level of sequence conservation at any given position.

**Figure 4.**
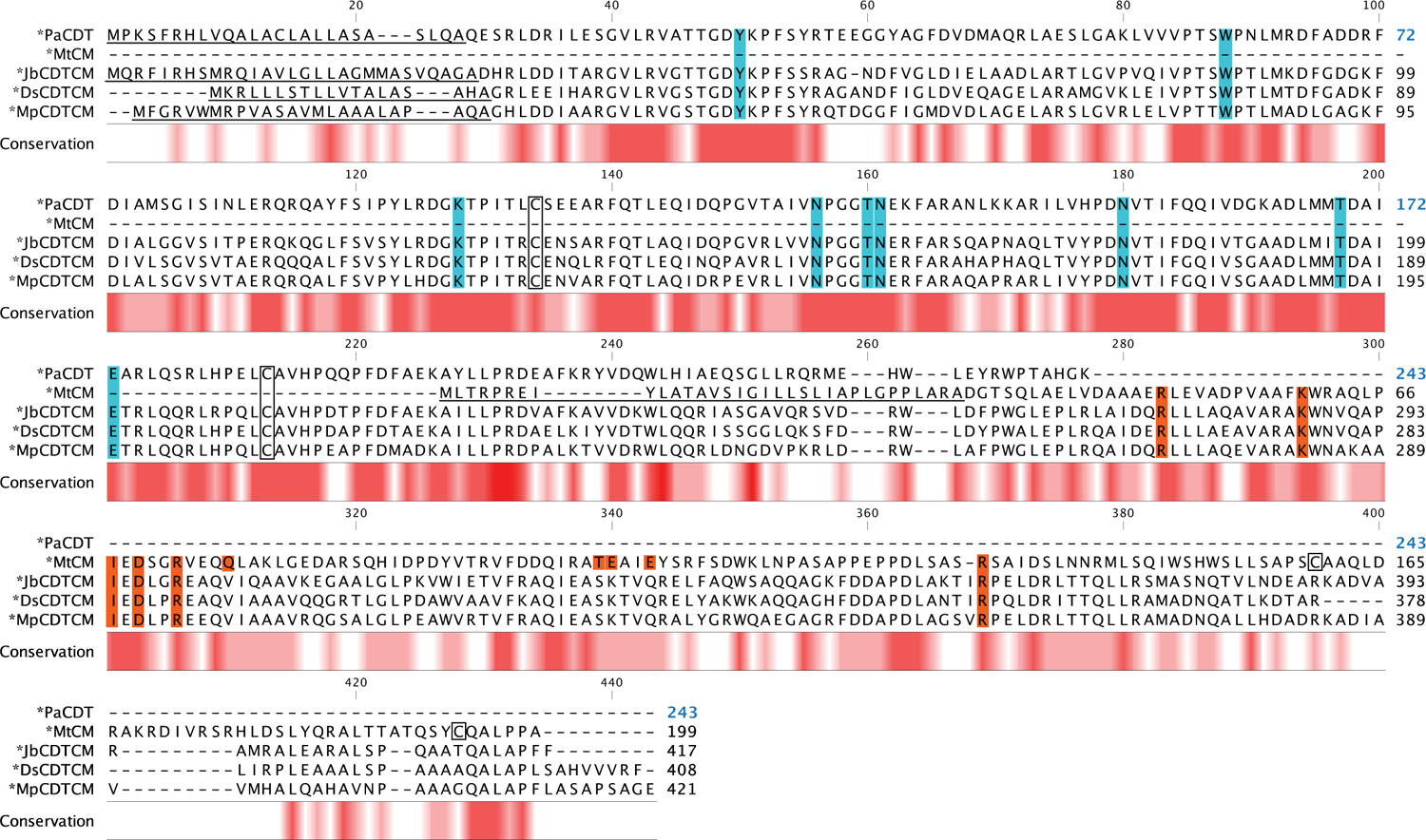
Sequence alignment of bifunctional *CDTCM variants. Shown is the amino acid sequence alignment of the bifunctional *CDTCM fusion enzymes to *MtCM and *PaCDT. More details are described in the legend to Figure 3.

Based on the known active site residues of *PaCDT^32^ and *MtCM,^9, 19^ the corresponding residues in the bifunctional fusion enzymes could be tentatively assigned. The nine active site residues of *PaCDT (Tyr22, Trp60, Lys100, Asn128, Thr132, Asn133, Asn152, Thr169, and Glu173)^32^ are fully conserved in all investigated exported bifunctional fusion enzymes, whereas of the 10 active site residues of *MtCM (Arg49, Lys60, Ile67, Asp69, Arg72, Gln76, Thr105, Glu106, Glu109, and Arg134),^9^ only 50% are conserved. Residues Gln76 and Glu109 are substituted in all bifunctional enzymes with Val and Gln, respectively. For residue Thr105, an Ala was found in all *CMCDTs and a Ser in all *CDTCMs. Glu106 is exchanged with a Lys in all bifunctional enzymes except for *ScCMCDT, where it is a Ser.

Additionally, the strictly conserved Cys residues forming a disulfide bond in *PaCDT are present in all CDT domains of the bifunctional fusion enzymes. In contrast, the two cysteines, which form a disulfide bond in *MtCM, are not conserved in the CM domains. The disulfide bond in *MtCM is located near the C-terminus in a segment that shows the highest sequence diversity among the bifunctional fusion enzymes (Figure 3 and Figure 4).

The genes of seven bifunctional fusion enzymes were custom-synthesized (Figure S2) and cloned into plasmids for expression in *E. coli* for further investigation. Of the *Shewanella* and *Thalassomonas* genera, only the variants from *S. baltica* and *T. actiniarum* were further analyzed.

### Complementation of CM and CDT-deficient mutant strains of *E. coli* by bifunctional fusion enzymes

The amino acid sequences of the four exported *CMCDTs (*ScCMCDT, *AfCMCDT, *TaCMCDT, and *SbCMCDT) and the three exported *CDTCMs (*JbCDTCM, *DsCDTCM, and *MpCDTCM), including their N-terminal natural signal peptides and an appended C-terminal His-tag, were reverse translated and codon-optimized for gene expression in *E. coli*. The custom-synthesized genes were cloned (without their signal sequence) into the expression vector pKTCTET-0^46^ for an initial assessment of heterologous CM and CDT activities *in vivo* inherent to the fusion proteins.

The *in vivo* complementation assay was carried out in *E. coli* strain KA12,^47^ which is devoid of the two endogenous genes encoding the bifunctional CM-prephenate dehydrogenase (PDH) and CM-prephenate dehydratase (PDT, a cyclohexadienyl dehydratase specific for prephenate), thus rendering KA12 auxotrophic for growth in minimal medium M9c^46^ that lacks Phe and Tyr. When KA12 is provided with the helper plasmid pKIMP-UAUC,^48^ which encodes a monofunctional PDH from *Erwinia herbicola* (*TyrA) and the CDT of *P. aeruginosa* (PheC, *PaCDT), only the CM activity is missing to restore growth in M9c medium. However, if KA12 is provided with the helper plasmid pKIMP-UA, which only encodes *TyrA, growth in M9c requires both an active CM and CDT.

Single colonies of KA12/pKIMP-UAUC and KA12/pKIMP-UA, additionally transformed with a pKTCTET expression plasmid that carries a gene for the fusion proteins were streaked out onto different M9c minimal agar plates. The plates contained different concentrations of tetracycline, the inducer of the promoter on pKTCTET, to probe for the strength of the *in vivo* complementation. Figure S3 shows that all bifunctional enzymes complemented the CM deficiency in KA12/pKIMP-UAUC cells, producing well visible colony growth within 4 days on M9c with 200 ng/mL tetracycline, except for *TaCMCDT that grows significantly slower than all other clones. Since the CM reaction precedes the CDT reaction, the growth in KA12/pKIMP-UA transformants is also dependent on the activity of the CM domain, and the observed growth phenotypes generally parallel those for the KA12/pKIMP-UAUC cells (Figure S3). Significantly lower complementation of the combined CM+CDT deficiency was found for *SbCMCDT at a low gene induction level. After 5 days at 37°C, all bifunctional fusion enzymes provided CM and CDT activity *in vivo* at least with 200 ng/mL tetracycline, except for *TaCMCDT lacking CDT activity.

### Bifunctional fusion enzymes show high catalytic efficiency *in vitro*

The catalytic activity of the bifunctional fusion enzymes was next investigated by kinetic activity measurements *in vitro*. The pKTCTET plasmids encoding the leaderless bifunctional fusion enzymes were transformed into the *E. coli* production strain KA29^19^ which harbors a chromosomal T7 RNA polymerase for high transcription levels from the T7 promoter on pKTCTET-0^46^ Besides lacking CM and PDT genes to exclude *a priori* contamination with endogenous enzyme activity, KA29 is deficient in the thioredoxin B (*trxB*) gene. This renders the cytoplasm more oxidative and facilitates disulfide bond formation, which is expected to occur in the CDT domain. Cytoplasmically overproduced bifunctional fusion enzymes were purified *via* their C-terminal His-tag using Ni^2+^-NTA affinity chromatography. All variants but *TaCMCDT could be produced in *E. coli* as soluble proteins. That *TaCMCDT remained insoluble may explain the impaired *in vivo* complementation in both KA12/pKIMP-UAUC and KA12/KIMP-UA by this fusion enzyme. Further *in vitro* kinetic studies were thus only performed with the six soluble bifunctional fusion enzymes. The bifunctional fusion enzymes were subjected to measurements of their isolated CM and CDT activities as well as the coupled CM+CDT kinetic activity. The derived Michaelis-Menten kinetic parameters are listed in Table 1.

**Table 1.**
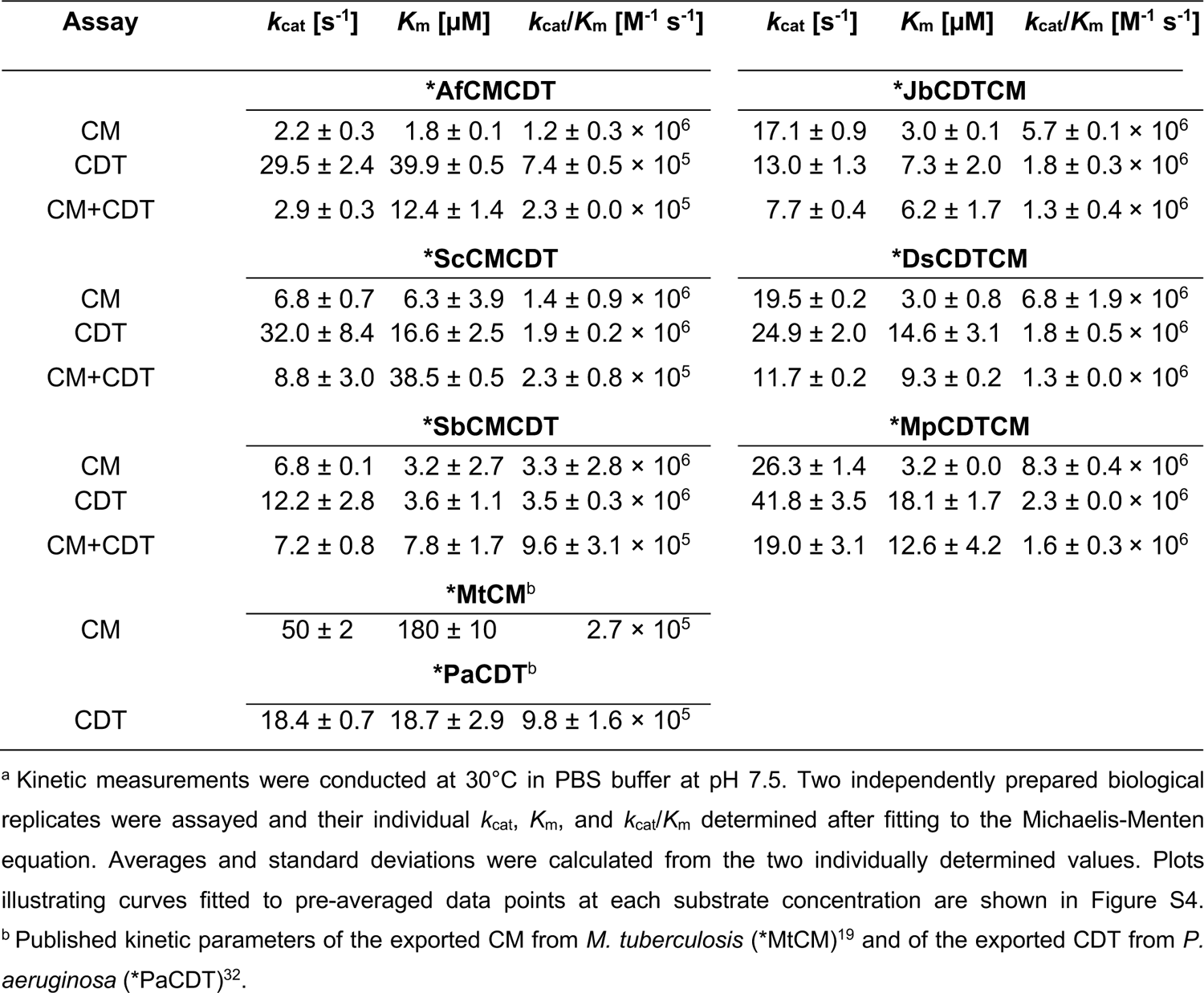
*In vitro* kinetic parameters of the bifunctional fusion enzymes^a^

All bifunctional fusion enzymes confirmed CM and CDT activities *in vitro*. Strikingly, all CM kinetics revealed a very low *K*_m_ <7 µM, which is at least 25-times lower than the *K*_m_ of 180 µM from *MtCM. Thus, the CM domains of the fusion enzymes can effectively bind very low concentrations of chorismate. The low *K*_m_ values are coupled to slower rate constants, with *k*_cat_ <7 s^-1^ or <26 s^-1^ determined for the *CMCDTs or the *CDTCMs, respectively, which is 2 – 25x lower than the *k*_cat_ of 50 s^-1^ reported for *MtCM.^19^ This suggests that the low *K*_m_ values may come at a cost. With the extraordinarily low *K*_m_ values, the catalytic efficiencies *k*_cat_/*K*_m_ of the CM domains from the bifunctional fusion enzymes reached 1 – 4 × 10^6^ M^-1^ s^-1^ for the *CMCDTs and 5 – 8 × 10^6^ M^-1^ s^-1^ for the *CDTCMs and are an order of magnitude higher than *k*_cat_/*K*_m_ of 2.7 × 10^5^ M^-1^ s^-1^ for *MtCM.^19^

The kinetic parameters of the CDT domains from all bifunctional enzymes are similar to those of the monofunctional reference enzyme *PaCDT. Individual rate constants and Michaelis constants for the bifunctional fusion enzymes vary from 12 – 42 s^-1^ and 3 – 40 µM, respectively, but generally are in the same order of magnitude as those of *PaCDT, with a *k*_cat_ of 18.4 s^-1^ and a *K*_m_ of 18.7 µM. All catalytic efficiencies lie around 10^6^ M^-1^ s^-1^, and no differences in CDT activity are apparent between the *CMCDT and *CDTCM variants.

The coupled CM+CDT kinetic measurement allowed first insights into whether the domain fusion influences the catalytic activity, as it simultaneously measures the two consecutive conversions from chorismate *via* prephenate to phenylpyruvate. In the absence of a domain fusion effect, the poorer of the individual kinetic parameters from both catalytic steps would limit the coupled reaction velocity and reveal the lower *k*_cat_ and higher *K*_m_ of either individual domain.

The results of the coupled CM+CDT assay for the *CMCDTs demonstrate that the rate constant (*k*_cat_) of the CM step was limiting, whereas the Michaelis constant (*K*_m_) was higher than for each of the individual steps for *ScCMCDT and *SbCMCDT. In contrast, *AfCMCDT exhibited a *K*_m_ of 12.4 µM for the coupled CM+CDT activity, which is lower than the measured *K*_m_ of 39.9 µM for CDT activity alone. This suggests that substrate saturation for the coupled CM+CDT reaction is reached at a lower substrate concentration. However, this effect could be explained by a lag or transient phase, which can occur in coupled enzyme reactions, that lasts until steady-state conditions for the downstream enzymes are reached.^49–52^

A similar observation was made for the *CDTCM variants. Also this inverse format did not reveal an increased catalytic rate constant *k*_cat_ for the coupled CM+CDT assay compared to the single CM or CDT assays. The *K*_m_ values of the coupled CM+CDT assays lie between those for the individual catalytic steps, like for *AfCMCDT. Remarkably, the coupled CM+CDT reactions exhibited lower *k*_cat_ values compared to the singly measured CM and CDT activities, suggesting that neither of the domains plays a dominant role for limiting catalysis. Possibly, a rate-limiting conformational change is required to release prephenate from the CM domain, such that it can rebind in the active site of the CDT domain, which would also explain the slower rate constant. In summary, the coupled CM+CDT catalytic efficiencies did not surpass the individually measured CM nor CDT activities.

Structural insight into the relative topologies of the catalytic centers of a fusion enzyme is essential to study direct interactions that may, for instance, facilitate the rebinding of prephenate released from the CM domain by the CDT domain. Close spatial interactions would be possible by either having the CM and CDT active sites next to each other within the same folded single polypeptide or upon head-to-tail dimerization of two bifunctional proteins. To address the latter, we investigated the quaternary structure of selected bifunctional fusion proteins in solution.

### Investigation of quaternary structures in solution

Homodimer formation was reported for *MtCM,^9^ and homotrimer formation for *PaCDT.^32^ To investigate whether the exported bifunctional fusion enzymes form oligomeric structures in solution, size-exclusion chromatography (SEC) was performed with all variants included in this investigation except for *SbCMCDT and *TaCMCDT as they did not yield sufficient soluble protein for SEC analysis.

All bifunctional enzymes eluted in a large prominent peak at around 15 mL that corresponds to the molecular mass of a monomer (Figure 5). In addition, a small peak was observed between the 150 kDa and 44.3 kDa markers of the protein standard, corresponding in mass to a dimer. This peak was detected for all *CDTCM variants (Figure 5B; red arrowhead) and possibly for *ScCMCDT, which displayed a small hump at this elution volume (Figure 5A). There is no indication of dimer formation of *AfCMCDT; however, we noted a right-hand shoulder of the main peak (Figure 5A), which could be due to conformational isomers, *e.g.*, due to partial unfolding. The wide shoulder eluting ahead of the 670 kDa protein standard for *AfCMCDT and *DsCDTCM (Figure 5A and Figure 5B) suggests significant protein aggregation in these samples. An aggregation propensity could also explain the observed small dimer peak for most of the bifunctional fusion enzymes.

**Figure 5.**
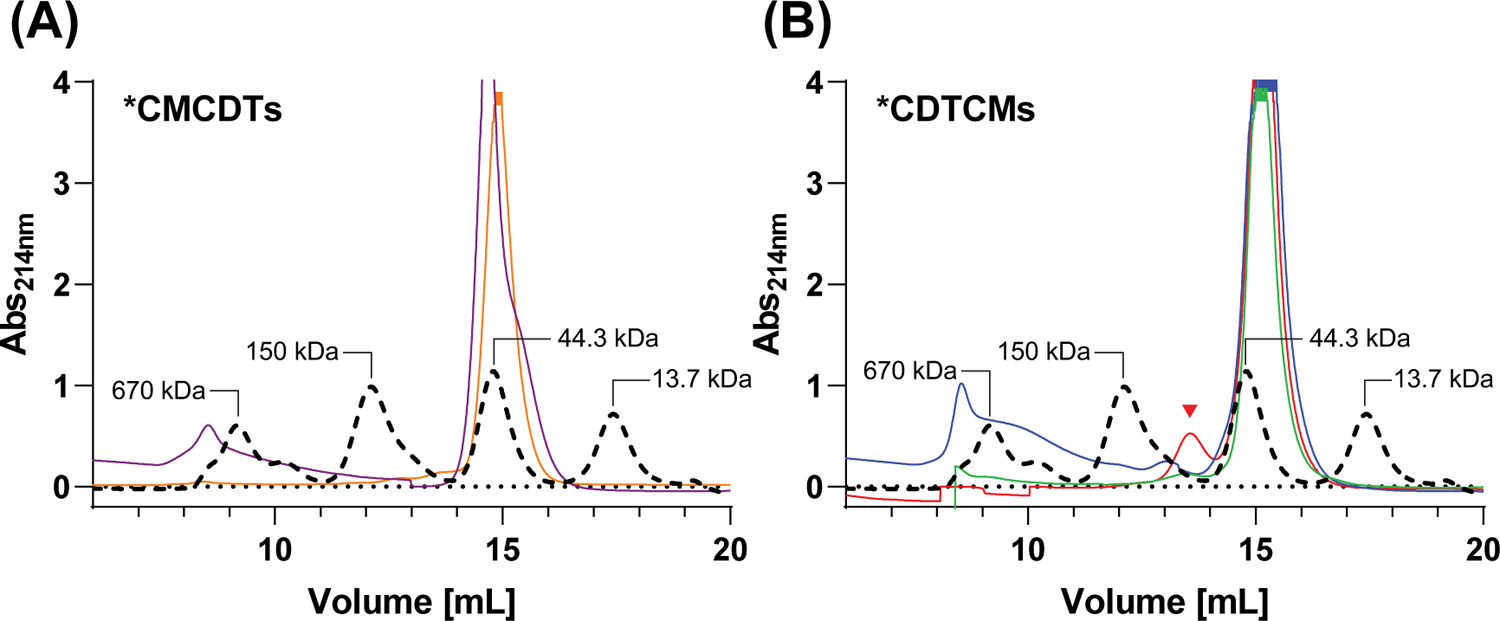
Quaternary structure analysis of bifunctional fusion enzymes. Shown are the SEC elution profiles at 214 nm absorbance. The standard protein mix (black dashed line, molecular masses indicated) is overlaid with the profiles of (A) *CMCDTs, including *AfCMCDT (purple) and *ScCMCDT (orange), and with (B) *CDTCMs, including *JbCDTCM (green), *DsCDTCM (blue), and *MpCDTCM (red). The red arrowhead indicates the peak of a possible dimer.

### Crystal structures of exported bifunctional fusion enzymes

To investigate the relative locations of the active sites within a single polypeptide, *AfCMCDT, *JbCDTCM, and *DsCDTCM were crystallized, and their structures solved and refined to 1.6, 1.7, and 2.4 Å resolution, respectively, and *R_free_*/*R_work_* values of 24.8/20.5%, 24.3/19.9%, and 25.8/23.0% (Table 2). We thus have high-quality examples for both *CMCDT and *CDTCM topologies, represented by *AfCMCDT and *JbCDTCM, respectively, and a lower-quality structure of *DsCDTCM, with relatively high Wilson *B*-factors indicating disorder (Table S1).

**Table 2.** Data collection and refinement statistics

|  | *AfCMCDT | *JbCDTCM, MES | *DsCDTCM |
| --- | --- | --- | --- |
| **Data collection** |  |  |  |
| Beamline/Synchrotron | ID30A-1/ESRF | BioMAX/MAX IV | BioMAX/MAX IV |
| Wavelength (Å) | 0.965 | 0.976 | 0.984 |
| Space group | $P2_1$ | $P2_12_12_1$ | $P2_12_12_1$ |
| Unit cell parameters |  |  |  |
| $a, b, c$ (Å) | 36.4 98.8 55.3 | 40.6, 74.0, 131.2 | 92.1, 111.9, 221.4 |
| $\alpha, \beta, \gamma$ (°) | 90 108.5 90 | 90 90 90 | 90 90 90 |
| Number of protein chains in a.u. | 1 | 1 | 4 |
| Resolution range<sup>a</sup> | 52.5 – 1.55 (1.73 – 1.55) | 65.6 – 1.65 (1.83 – 1.65) | 99.9 – 2.44 (2.66 – 2.44) |
| Total reflections<sup>a</sup> | 217488 (15214) | 445335 (17372) | 685834 (33367) |
| Unique reflections<sup>a</sup> | 33225 (2216) | 35076 (1754) | 51859 (2594) |
| Spherical completeness<sup>a</sup> (%) | 61.7 (14.7) | 72.1 (13.5) | 60.3 (13.5) |
| Ellipsoidal completeness<sup>a</sup> (%) | 88.9 (69.7) | 93.0 (54.6) | 94.0 (72.7) |
| CC<sub>1/2</sub><sup>a</sup> | 99.7 (52.9) | 99.8 (54.2) | 99.4 (12.4) |
| Multiplicity<sup>a</sup> | 6.5 (6.9) | 12.7 (9.9) | 13.2 (12.9) |
| Mean $I/\sigma(I)$<sup>a</sup> | 9.3 (1.6) | 14.1 (1.6) | 7.8 (1.5) |
| **Refinement** |  |  |  |
| Resolution range (Å) | 52.5 – 1.55 | 65.6 – 1.65 | 99.9 – 2.44 |
| $R_{free}/R_{work}$ (%)<sup>b</sup> | 24.8 / 20.5 | 24.3 / 19.9 | 25.8 / 23.0 |
| Average $B$-factors (Å<sup>2</sup>) | 26.2 | 30.3 | 59.5 |
| Number of non-hydrogen atoms | 3269 | 3587 | 12318 |
| Protein | 3139 | 3421 | 12234 |
| Water | 117 | 154 | 71 |
| Ligands | 13 | 12 | 10 |
| Protein residues | 389 | 393 | 1542 |
| r.m.s.d. from ideal geometry |  |  |  |
| Bond lengths (Å) | 0.010 | 0.010 | 0.007 |
| Bond angles (deg.) | 1.64 | 1.74 | 1.11 |
| Ramachandran plot |  |  |  |
| Favored (%) | 95.8 | 97.7 | 96.4 |
| Outliers (%) | 0.3 | 0.0 | 0.3 |
| PDB ID | 8CQ3 | 8CQ4 | 8CQ6 |

<sup>a</sup> Values in parentheses refer to highest resolution shell.
<sup>b</sup> $R_{free}$ was calculated from 5% of randomly selected reflections for each data set.

Superimposition of *MtCM (Protein Data Bank (PDB) ID: 2FP2)^9^ and *PaCDT (PDB ID: 5HPQ)^32^ structures onto the CM and CDT domains of *AfCMCDT (PDB ID: 8CQ3; this work), *JbCDTCM (PDB ID: 8CQ4; this work) and *DsCDTCM (PDB ID: 8CQ6; this work) revealed highly similar structures in the bifunctional fusion enzymes (Figure 6). Like *MtCM, the CM domains of *AfCMCDT, *JbCDTCM, and *DsCDTCM form six-helix bundles with a single active site in the N-terminal moiety. The CDT domains comprise two α/β subdomains that are connected by two antiparallel hinge β-strands, crossing between the sub-domains twice, as known for *PaCDT.^32^ Also the structures of *DsCDTCM and *JbCDTCM are very similar, both overall and in the active site (Figure S5), with r.m.s.d. = 1.4 Å (C_α_ atoms). We noticed that *JbCDTCM contained a 2-(*N*-morpholino)ethanesulfonic acid (MES) molecule from the buffer in its CDT active site (Figure 6, Figure S5C, and Figure S6C). *AfCMCDT captured acetate molecules from the crystallization buffer, in both the CM and CDT active sites (Figure S6A and S6B).

**Figure 6.**
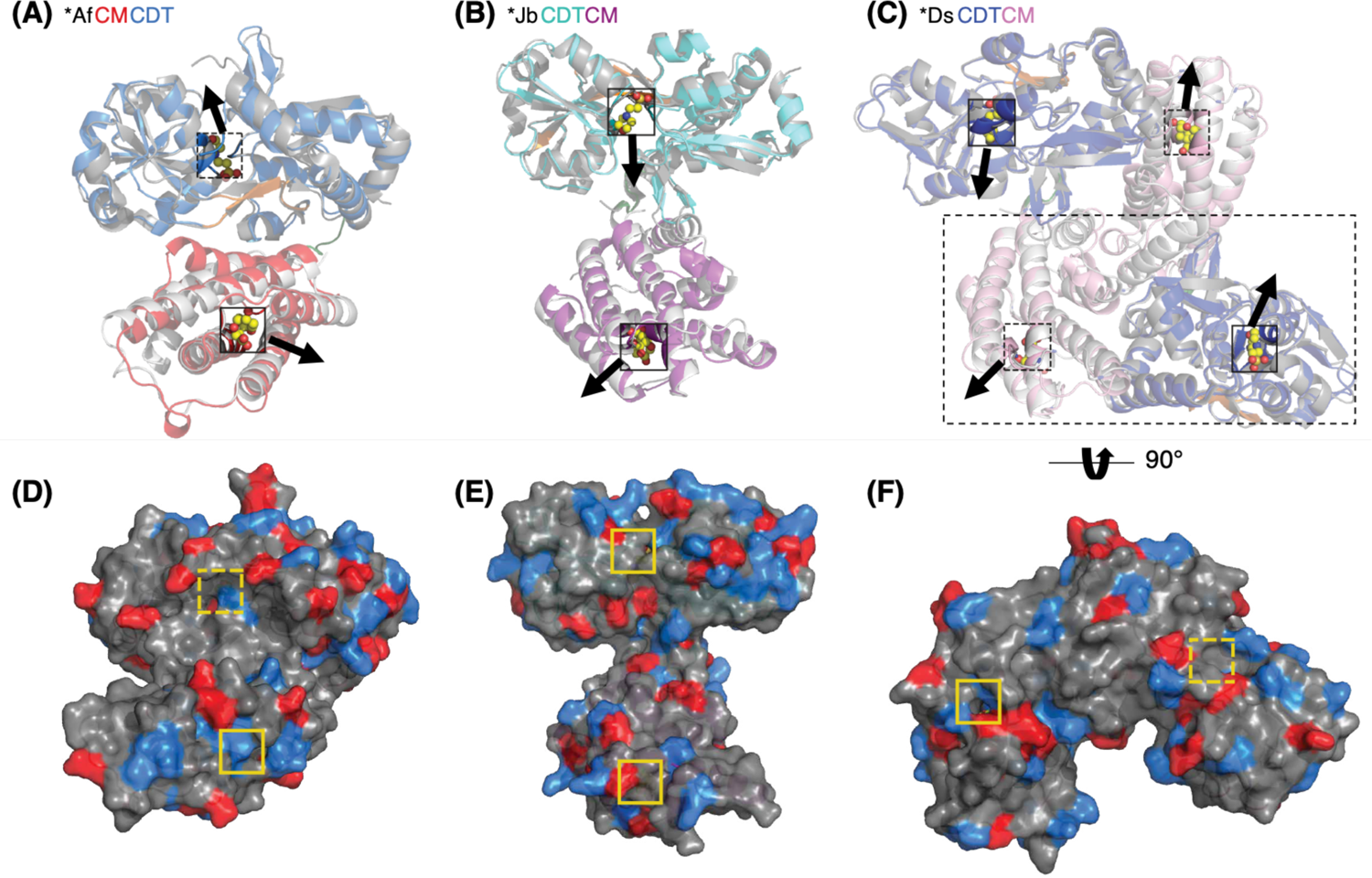
Superimpositions of *AfCMCDT, *JbCDTCM, and *DsCDTCM with *MtCM and *PaCDT. (A-C) *MtCM (PDB ID: 2FP2, light grey)^9^ and *PaCDT (PDB ID: 5HPQ, dark grey)^32^ are shown superimposed onto *AfCMCDT (PDB ID: 8CQ3; this work) (A), *JbCDTCM (PDB ID: 8CQ4; this work) (B), and *DsCDTCM dimer (PDB ID: 8CQ6; this work) (C), with CM and CDT domains from two different chains shown in a large dashed rectangle. CM domains of the fusion proteins are depicted in red, magenta, and pink, respectively, and CDT domains in blue, cyan, and dark blue. Hinges between CDT subdomains are colored orange, and the expected exit trajectories from the active sites are indicated with black arrows. (D-F) Surface representations of *AfCMCDT (PDB ID: 8CQ3; this work) (D), *JbCDTCM (PDB ID: 8CQ4; this work) (E), and *DsCDTCM (PDB ID: 8CQ6; this work) (F) with residues charged positively (Arg, Lys, His) and negatively (Glu, Asp) in blue and red, respectively. Note that the domains in the dashed rectangular box are the CM and CDT domains from the two different chains shown in (C), but rotated by 90° in panel (F) for better surface visualization. The active sites are indicated with small squares; those facing backwards drawn with dashed lines. The CM active sites are illustrated with the *endo*-oxabicyclic transition state analog (TSA) bound in the *MtCM structure (PDB ID: 2FP2)^9^, and the CDT active sites with a 2-(*N*-morpholino)ethanesulfonic acid (MES) molecule that was co-crystallized with *JbCDTCM (yellow spheres; for details, see Fig. S5C).

### Active sites

*JbCDTCM and *DsCDTCM share the same residues in both CM and CDT active sites (Figure S5B and S5C). Therefore only *JbCDTCM will be discussed as representative of the *CDTCM topology. Six of ten active site residues of *MtCM are conserved in the CM domains of *AfCMCDT and *JbCDTCM (Figure 7A).^9^ Compared to *MtCM, the active site residues in *CDTCM and *CMCDT fusion enzymes differ at several positions, *e.g.*, Q76V and T105S or T105A (in *JbCDTCM or *AfCMCDT, respectively), E106K, and E109Q (Figure 7A). In *MtCM, Gln76 and Thr105 help to coordinate the ligand’s carboxylates,^9^ which is no longer possible upon substitution with Val or Ala as in *AfCDTCM, whereas the substitution of Thr105 by a Ser in *JbCMCDT is more conservative and could still provide a hydrogen bond donor to the carboxylate *via* its hydroxyl group. Likewise, substituting Glu106 with Lys could, together with some conformational adaptation of the side chain, preserve coordination of the ligand’s hydroxyl group (shown for *MtCM),^9^ and substituting Glu109 with Gln could accomplish coordination of the ether oxygen *via* its amide hydrogen instead of the carboxyl group. However, by replacing Glu106 and Glu109 with positively charged or neutral residues, the charge balance of the active site is affected. Importantly, this likely explains why the fusion enzymes exhibit lower rate constants *k*_cat_ and increased substrate affinity (lower *K*_m_) in the CM kinetic assays, compared to *MtCM (Table 1).

**Figure 7.**
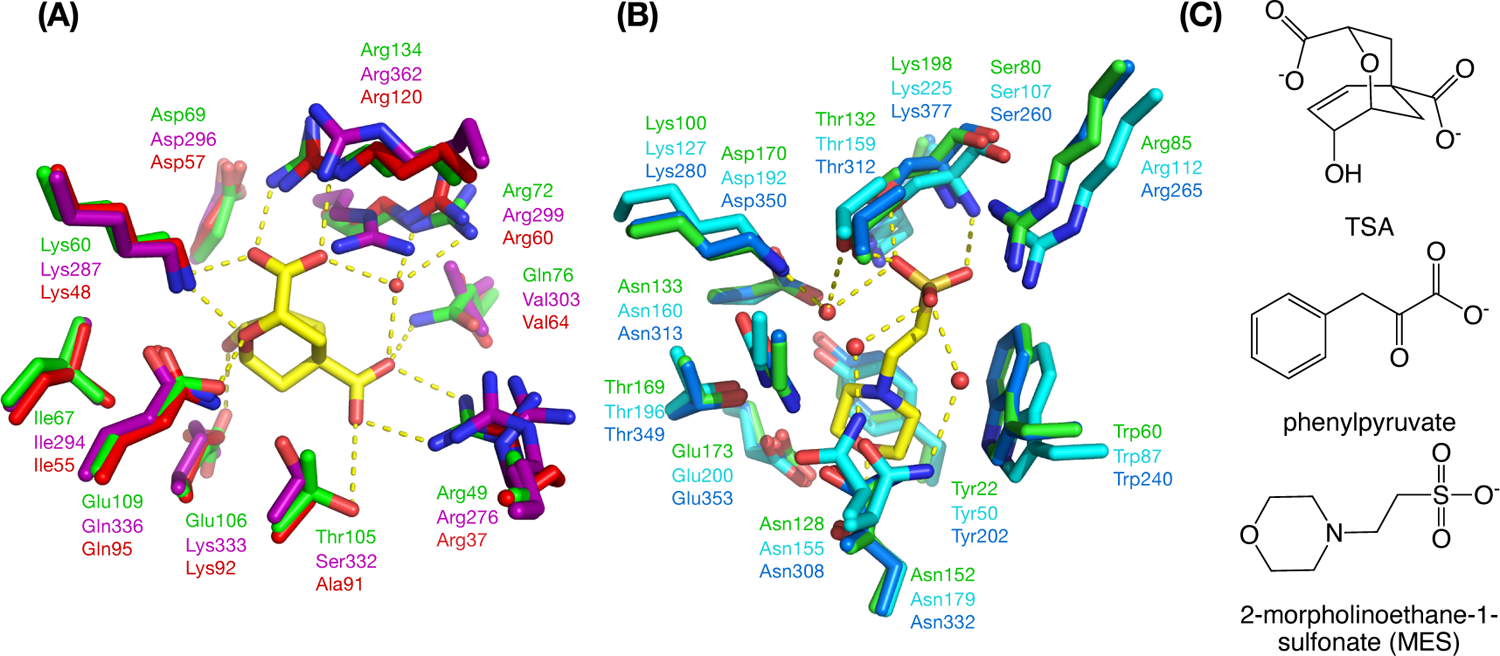
Superimposition of active site residues in CM and CDT. (A) Superimposition of CM active sites of *AfCMCDT (PDB ID: 8CQ3; this work) (red) and *JbCDTCM (PDB ID: 8CQ4; this work) (magenta) with the previously published monofunctional AroQγ CM holotype *MtCM (PDB ID: 2FP2, green)^9^ in complex with an *endo*-oxabicyclic transition state analog (TSA, yellow; CM active sites of *AfCMCDT and *JbCDTCM are not occupied). (B) Superimposition of CDT active sites of *AfCMCDT (PDB ID: 8CQ3; this work) (blue) and *JbCDTCM (PDB ID: 8CQ4; this work) (cyan) with the monofunctional *PaCDT (PDB ID: 5HPQ, green)^32^. *JbCDTCM contains a MES molecule from the buffer (yellow) in the CDT active site. (C) Chemical structures of the ligands TSA, phenylpyruvate (the CDT substrate), and MES. Hydrogen-bonding interactions between ligand and protein are depicted as yellow dashed lines. Water molecules are displayed as red spheres.

In contrast to the analyzed CM domains, all nine residues forming the active site in *PaCDT^32^ are fully conserved not only in the CDT domains of *AfCMCDT and *JbCDTCM (Figure 7B), but in all investigated exported bifunctional fusion enzymes (Figure 3 and Figure 4). The CDT domains of the three bifunctional enzymes show very high structural similarity to *PaCDT, and congruently the determined CDT kinetic parameters for *AfCMCDT and *JbCDTCM are highly similar to those of *PaCDT (Table 1).

In the active site of the *JbCDTCM structure, we discovered a MES molecule (Figure S6C). MES resembles the CDT product, phenylpyruvate (Figure 7C), and thus allows for the first glimpse into the enzyme-substrate interactions. The carboxylate group of phenylpyruvate is replaced by a sulfonate group in MES, and the aromatic ring by a morpholine ring (Figure 7C). The sulfonate group of MES forms hydrogen bonds with the hydroxyl group and backbone amide of Ser107 and Thr159 in *JbCDTCM and interacts by charge complementarity with the side chains of Arg112 and Lys225 (Figure 7B and Figure S5C). Water-mediated protein interactions of the CDT domain with the MES sulfonate group involve Lys127, Asp192, Asn155, and Thr159. The morpholine ring is stabilized by interactions with the side chains of Tyr50 and Trp87. The oxygen atom in the morpholine ring is hydrogen-bonded to the side chain amide nitrogen of Asn179 (Figure 7B and Figure S5C).

### Structural evidence for substrate gating and channeling

Since fused enzymes often exhibit cooperation between active sites, we were interested in the relative placements of these. In *AfCMCDT, representing the *CMCDT topology, the presumed substrate access to the CDT domain faces away from the expected product exit site of the CM domain^9^ (Figure 6A), with a distance of 35 Å between active sites. In the two *CDTCMs, the distances between CM and CDT active sites are similarly large, measuring approximately 40 Å, thus disfavoring a direct intramolecular transfer of the metabolic intermediate prephenate from the CM to the CDT active site. Surface representations of the bifunctional enzymes show accessible CM active sites for all three enzymes regardless of the ligand, while the CDT active site is solvent-accessible only in MES-bound *JbCDTCM.

Whereas *AfCMCDT and *JbCDTCM crystals contained only one polypeptide chain in the asymmetric unit, the *DsCDTCM crystal contained four chains; with head-to-tail dimers (CM-to-CDT) established by crystallographic symmetry (Figure 6C, Figure S7A and S7B). The buried surface area of this *DsCDTCM dimer was estimated to be 3070 Å^2^, with the free energy of dimer dissociation being 5.8 kcal/mol, as calculated using PDBePISA.^53^ The dimerization interface is mainly lined with hydrophilic and charged residues (Figure S7C and S7D). Small humps reminiscent of *CDTCM dimers in size-exclusion chromatography (Figure 5B) could be explained by transient formation of such dimers in the protein preparations.

The crystal structure of *DsCDTCM head-to-tail dimers (Figure 6C) allowed us to address possible intermolecular substrate channeling between domains of different protomers in a protein complex. The active sites of the CM and CDT domains of the two individual chains are separated by a significant distance of about 40 Å and open towards opposite directions (Figure 6C and 6F). The surface charge distribution (Figure 6D-F) does not reveal any pattern for a potential intra or inter-molecular electrostatic relay of intermediates from the CM to the CDT active site. Preliminary Molecular Dynamics simulations performed with the crystal structures of *AfCMCDT and *JbCDTCM (not shown) also gave no indication for substrate channeling. Taken together, the crystal structures do not provide evidence for intra or inter-molecular substrate channeling for the bifunctional fusion enzymes investigated here.

### Interactions at the CM and CDT domain interface

Whereas CDT domain superimpositions with *PaCDT revealed no prominent structural differences (Figure 6A-C), the two C-terminal helices H5 and H6 of the CM domains in *CMCDT and *CDTCM align with a slightly different angle, with *CDTCMs more closely matching *MtCM (Figure 8A, exemplified with *JbCDTCM). This suggests that C-terminal fusion to CDT may affect the CM structure. In *MtCM, a disulfide bond connects these two C-terminal helices, and it was pointed out that this crosslinking is important for the enzyme’s stability outside the cell.^54^

**Figure 8.**
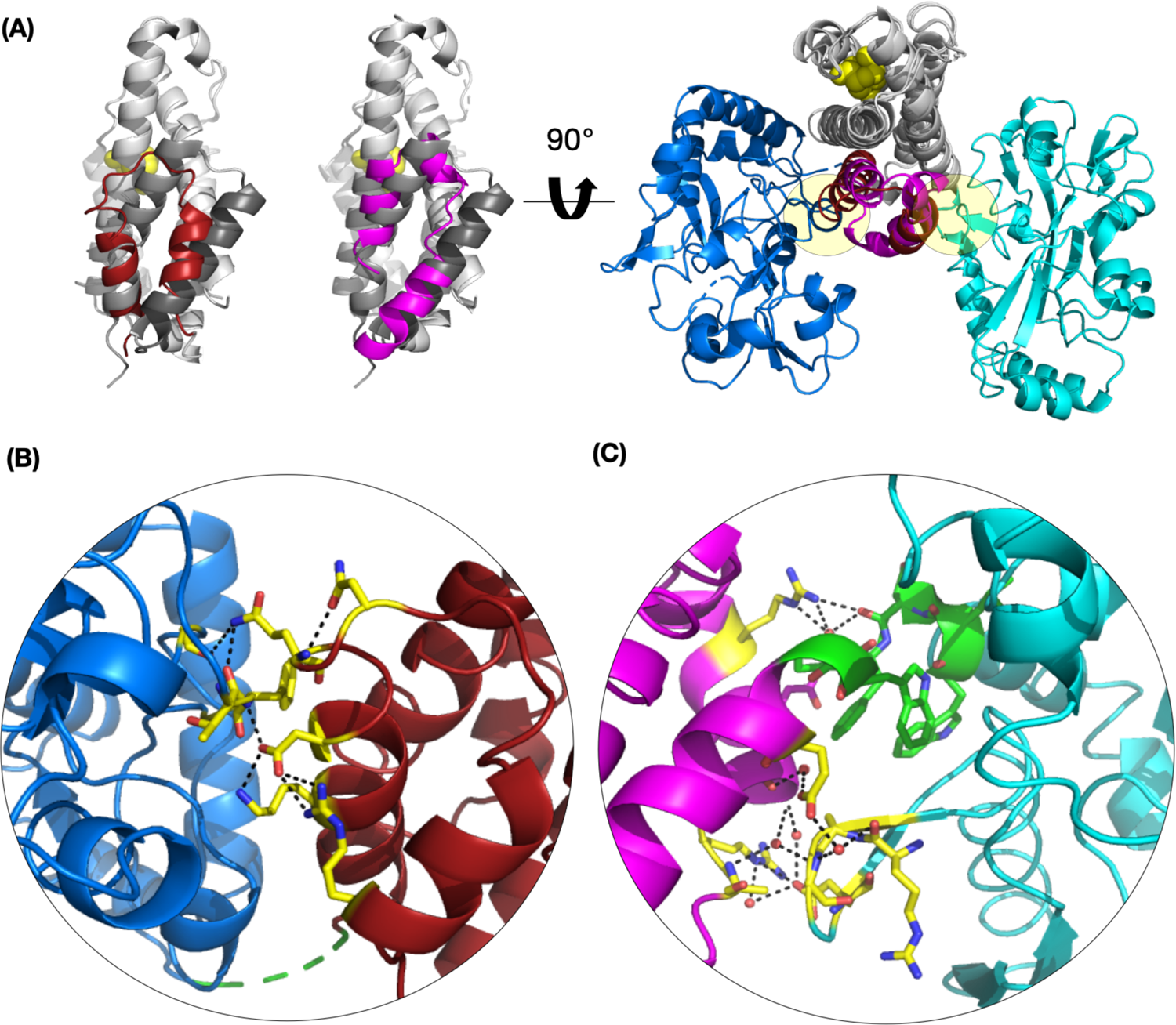
Protein-protein interactions between CM and CDT fusion domains. (A) Left panel: Superimposition of *MtCM (PDB ID: 2FP2)^9^ (white, with helices 5 and 6 in gray) with the CM domains of *AfCMCDT and *JbCDTCM (respective PDB IDs: 8CQ3 and 8CQ4; this work; both in white, with their corresponding C-terminal helices in red and magenta, respectively). The TSA ligand (yellow spheres) indicates the position of CM active site pockets relative to helices 5 and 6. Right panel: CM domains of *AfCMCDT and *JbCDTCM, superimposed to illustrate the topological difference of their corresponding CDT domains (blue and cyan, respectively), with the CM domains rotated 90° relative to the structures on the left. The circular lenses in (B) and (C) allow close-up views of the molecular interaction network between the CM and CDT domains of (B) *AfCMCDT and (C) *JbCDTCM (respective PDB IDs: 8CQ3 and 8CQ4; this work). The linker sequences (green) connect the CM and CDT domains; residues engaged in prominent polar contacts (yellow) between the two domains are shown as sticks. Water molecules involved in the interaction network are depicted as red spheres, and hydrogen bond contacts with black dashed lines. For electron density of the linker regions, see Figure S8.

A full structural assessment of the domain interface in *AfCMCDT is precluded by the poorly resolved linker region (Figure 8B). In fact, the Pro156-Asp159 stretch showed ambiguous electron density (Figure S8A). This suggests a more flexible arrangement of the CM and CDT domains for *AfCMCDT than for the topological counterparts *JbCDTCM and *DsCDTCM, for which the linker regions are clearly resolved (Figure S8B and S8C). In *JbCDTCM, CM helices H1 and H6 interact with the conserved ‘β-sheet extrusion’ of CDT (Figure 8C). This structural interaction in *CDTCMs appears to be fairly rigid, as the linker segment contains many aromatic and hydrophobic residues that are shielded from the aqueous environment (Figure 8C). Furthermore, the hydrogen bonding network between CM helix H5 and the β-sheet extrusion from the CDT domain shields the hydrophobic linker and forms fold-stabilizing hydrogen bonds with water molecules. It thus appears that the rigid domain interface in *CDTCMs assumes a stabilizing role for the secreted fusion enzyme, thereby taking over the function of the stabilizing disulfide bonds previously observed in secreted CMs.^9,19^

### Kinetic evidence for substrate gating and channeling from designed active site knock-out and split-domain variants

To shed light on potential functional linkages between the two catalytic sites, we designed *(i)* two-domain protein variants having knocked-out (KO) individual active sites and *(ii)* truncated variants with individual CM or CDT domains split away. Increased CDT activity in a CM active site KO variant (CDT or CDT) in comparison with the parental enzyme would suggest that the presence of an active CM domain interferes with CDT activity. Conversely, a CDT active site KO variant (CM or CM) may reveal potential effects of the CDT domain on CM domain function. Split variants consisting only of a single CM or CDT domain, thus uncoupling the enzymatic activities, may provide complementary information on the biological purpose of the domain fusions. *AfCMCDT and *JbCDTCM served as representatives of the two bifunctional fusion enzyme topologies for the generation of the active site KO and split domain variants.

The active site KO variants were generated by introducing a single residue substitution in either the CM or the CDT domain. In *E. coli* PheA (EcCM), Lys39 was reported to be essential for CM activity,^7, 55^ which was confirmed by a K39A substitution variant that showed 10^4^-10^5^-fold lower *k*_cat_/*K*_m_.^56^ Lys39 in EcCM corresponds to Lys60 in *MtCM,^54^ which was used to allocate the corresponding catalytic residue for substitution with Ala in *AfCDT (K48A) and *JbCDT (K287A) based on amino acid sequence alignments (Figure 3 and Figure 4). Glu173 was shown to be essential for CDT activity in *PaCDT, serving as the proton donor to the departing hydroxyl group of prephenate, and a substitution with Gln leading to complete inactivation of the dehydratase activity.^32^ Amino acid sequence alignments (Figure 3 and Figure 4) were consulted to design the corresponding Glu substitution with Gln in *AfCM(E353Q) and *JbCM (E200Q).

To decide on the split site for engineering the single domain variants, the crystal structures of *AfCMCDT and *JbCDTCM (Figure 8 and Figure S8) were analyzed in the context of the amino acid sequence alignment of all available natural fusion enzymes (Figure 9).

**Figure 9.**
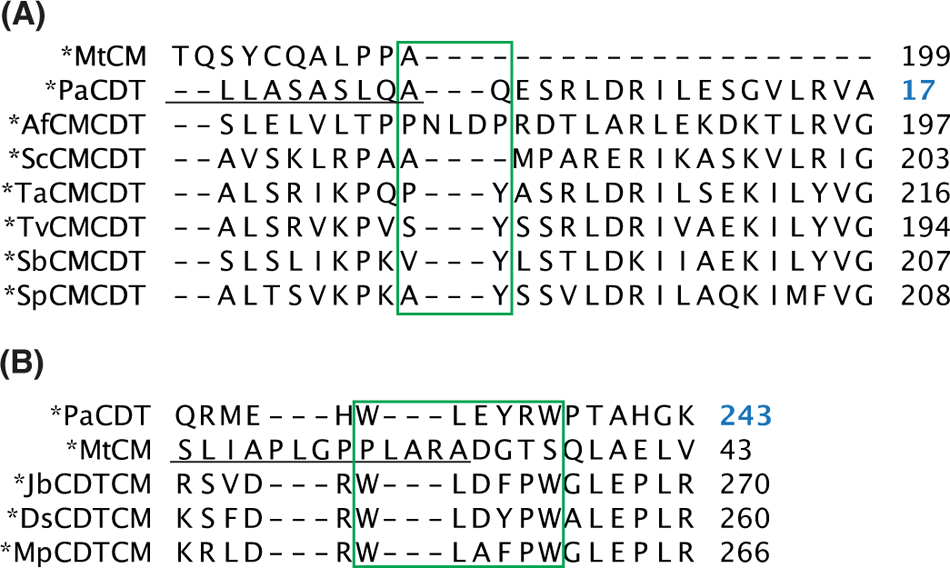
Amino acid sequence alignments corresponding to the domain linker regions. Underlined are amino acid residues belonging to the signal peptides. Residues that are presumed to be part of the linkers are identified with the green box. The numbers on the right refer to the right-most residue. Alignment of (A) *CMCDT and (B) *CDTCM variants.

For *AfCMCDT, the domain-linking segment could not be resolved (Figure S8A); therefore, the choice of the split site was based solely on the sequence alignment of Figure 9A. Since *AfCMCDT was the only *CMCDT variant with an elongated linker segment, no conserved feature or motif could be determined. Ultimately, the split site was placed between Leu179 and Asp180, in the middle of the linker sequence. For the *CDTCM variants the segment linking the two domains showed a highly conserved Trp-Leu-Xaa3-Xaa4-Xaa5-Trp (WLXXXW) motif, where Xaa3 frequently contributes a negative charge (Asp or Glu), Xaa4 is always Phe or Tyr, and Xaa5 is usually Pro. Residue Xaa5 also forms the start of the first α-helix in the CM domain (Figure 9B). Because the crystal structure of *PaCDT (PDB ID: 5HPQ)^32^ could only be resolved to Leu233 (just C-terminal to the first Trp in the linker segment), no structural information could be obtained from *PaCDT to help determine the split site for fully preserving the intact CDT domain. In the *JbCDTCM and *DsCDTCM structures presented here, we observed well-defined electron density for the linkers, with interactions of the two Trp residues at both ends (Figure S8B and S8C). Ultimately, the split site after the CDT domain was positioned at the end of the linker sequence, between Trp237 and Gly238, based on our new structural insights and the fact that the whole linker segment is conserved in all aligned CDTs. All active site KO variants as well as *AfCDT and *JbCM single domains were produced with a C-terminal His-tag, whereas *AfCM and *JbCDT single domains were cloned with an N-terminal His-tag. Expression of the reengineered genes in *E. coli* strain KA29 resulted in good soluble protein yields of all constructs, except for *AfCDT (little soluble protein) and *JbCDT (no soluble protein).

The CM activities of *JbCM *JbCM, and *AfCM match well with the corresponding catalytic parameters of the wild-type parent enzymes (Table 3 and Figure S9A and S9B). Whereas the kinetic curves of all variants derived from *JbCDTCM fully overlap, the split-off *AfCM domain showed a slightly higher *k*_cat_ (2.9 s^-1^) and a twofold higher *K*_m_ (4.5 µM) than wild-type *AfCMCDT (*k*_cat_ = 2.2 s^-1^, *K*_m_ = 1.8 µM). However, because of the limited sensitivity of the employed discontinuous kinetic assay in the low substrate concentration range, the observed small differences compared to *AfCMCDT are probably insignificant. A slight structural perturbation caused by removing the CDT domain might also explain the somewhat lower *k*_cat_/*K*_m_ of *AfCM. Knocking out the anticipated active site Lys in *AfCDT (K48A) and *JbCDT (K287A) fully abolished CM activity, providing experimental evidence for the predicted active site mechanism (Table 3 and Figure S9A and S9B).

**Table 3.** *In vitro* kinetic activities of active site KO and split domain variants of *AfCMCDT and *JbCDTCM^a^

|  | CM assay |  |  | CDT assay |  |  |
| --- | --- | --- | --- | --- | --- | --- |
| Variants | $k_{cat}$ [s<sup>-1</sup>] | $K_m$ [μM] | $k_{cat}/K_m$ [M<sup>-1</sup> s<sup>-1</sup>] | $k_{cat}$ [s<sup>-1</sup>] | $K_m$ [μM] | $k_{cat}/K_m$ [M<sup>-1</sup> s<sup>-1</sup>] |
| *AfCMCDT<sup>b</sup> | 2.2 ± 0.3 | 1.8 ± 0.1 | 1.2 ± 0.3 × 10<sup>6</sup> | 29.5 ± 2.4 | 39.9 ± 0.5 | 7.4 ± 0.5 × 10<sup>5</sup> |
| *AfCM~~CDT~~ | 1.9 ± 0.1 | 1.7 ± 0.3 | 1.1 ± 0.2 × 10<sup>6</sup> | No activity detected<sup>c</sup> |  |  |
| *Af~~CM~~CDT | No activity detected<sup>c</sup> |  |  | 33.3 ± 2.9 | 17.9 ± 0.7 | 1.9 ± 0.2 × 10<sup>6</sup> |
| *AfCM | 2.9 ± 0.5 | 4.5 ± 0.4 | 6.5 ± 0.5 × 10<sup>5</sup> | Not measured<sup>d</sup> |  |  |
| *AfCDT | Not measured<sup>d</sup> |  |  | 20.2 ± 4.1 | 28.2 ± 1.6 | 7.1 ± 1.0 × 10<sup>5</sup> |
| *JbCDTCM<sup>b</sup> | 17.1 ± 0.9 | 3.0 ± 0.1 | 5.7 ± 0.1 × 10<sup>6</sup> | 13.0 ± 1.3 | 7.3 2.0 | 1.8 0.3 × 10<sup>6</sup> |
| *Jb~~CDT~~CM | 15.6 ± 0.4 | 2.4 ± 1.3 | 7.9 ± 4.5 × 10<sup>6</sup> | No activity detected<sup>c</sup> |  |  |
| *JbCDT~~CM~~ | No activity detected<sup>c</sup> |  |  | 13.9 ± 0.3 | 4.2 ± 0.4 | 3.3 ± 0.2 × 10<sup>6</sup> |
| *JbCM | 16.4 ± 1.5 | 2.2 ± 0.3 | 7.6 ± 1.7 × 10<sup>6</sup> | Not measured<sup>d</sup> |  |  |

<sup>a</sup> All kinetic measurements were performed in PBS buffer, pH 7.5, at 30°C. The reported $k_{cat}$ , $K_m$ , and $k_{cat}/K_m$ parameters represent the mean and standard deviation of the parameters calculated for two independently prepared biological replicates, for which the kinetic data were individually fitted to the Michaelis-Menten equation. In Figure S9, corresponding plots are shown, however, with the curve fit performed through the mean activity at each substrate concentration.
<sup>b</sup> The kinetic parameters of wild-type \*AfCMCDT and \*JbCDTCM are taken from Table 1 for comparison.
<sup>c</sup> The engineered active site KO variants showed no measurable enzymatic activity for their mutated domain function.
<sup>d</sup> The split domain variants were only measured for their domain's inherent activity.

*AfCDT and *JbCDT exhibit identical *k*_cat_ values for their CDT activities as their wild-type parents (Table 3 and Figure S9C and S9D). Interestingly, the *K*_m_ determined for *AfCDT (17.9 µM) and *JbCDT (4.2 µM) was approximately twofold lower than for *AfCMCDT (39.9 µM) and *JbCDTCM (7.3 µM), respectively (Table 3), leading to a twofold higher catalytic efficiency *k*_cat_/*K*_m_. This may indicate that the native CM domain provides a competing binding site for the substrate prephenate, which would lower the local substrate concentration for the CDT reaction, showing up as a comparatively lower apparent CDT activity of the wild-type formats.

Finally, the split domain *AfCDT exhibited lower *k*_cat_ (20.2 s^-1^) and *K*_m_ (28.2 µM) values, but eventually the same catalytic efficiency as its wild-type counterpart (Table 3). However, as for *AfCM, structural perturbance due to the missing protein domain could be the cause for the observed change in the kinetic parameters of *AfCDT. Notably, no activity was measurable anymore after exchanging the presumed active site Glu353 and Glu200 with Gln in *AfCMand *JbCM, respectively, thus experimentally confirming the crucial importance of this active site residue for CDT activity (Table 3 and Figure S9C and S9D).

Next, we compared the efficiency of the two-step catalytic sequence from chorismate to phenylpyruvate that is observable upon combining individual active site KO or split domain variants of *AfCMCDT with that of the intact bifunctional fusion protein (Figure S10). The coupled CM+CDT reactions showed that equimolar concentrations of the active site KO variants *AfCM and *AfCDT complemented each other perfectly and reached a catalytic activity (*k*_cat_ = 2.8 ± 0.2 s^-1^, *K*_m_ = 9.2 ± 1.5 µM, *k*_cat_/*K*_m_ = 3.0 ± 0.7 × 10^5^ M^-1^ s^-1^) that coincides, within experimental error, with that of wild-type *AfCMCDT (Table 1). This result directly suggests that the enzyme fusion does not provide a catalytic benefit. An experiment mixing equimolar split *AfCM and *AfCDT domains revealed a lower CM+CDT activity (*k*_cat_ = 1.3 ± 0.2 s^-1^, *K*_m_ = 20.8 ± 7.7 µM, *k*_cat_/*K*_m_ = 6.9 ± 3.6 × 10^4^ M^-1^ s^-1^). In this case, the reduced activity is owed, in part, to the two-fold lower CM activity measured for the individual split *AfCM domain after cleaving off its fusion partner (Table 3), thereby possibly perturbing the structure of the active site. The inability to produce the soluble split *JbCDT domain precluded a similar set of measurements with the *JbCDTCM system.

### Probing for feedback regulation of *AfCMCDT and *JbCDTCM by phenylalanine or tyrosine

Intracellular CMs involved in biosynthetic pathways are known to be subject to feedback regulation^1–3^ involving various features such as additional allosteric domains,^57^ dynamic dimer interfaces,^16^ or complexes with partner enzymes, which enable sophisticated inter-enzyme allostery.^10, 17, 18, 58–62^ In contrast, no feedback inhibition has been detected for the exported enzymes *MtCM^19^ or *PaCDT.^63^ To test whether *AfCMCDT and *JbCDTCM are subject to feedback control, a coupled CM+CDT assay was performed in the presence of a large excess of either Phe or Tyr, the relevant end products of the shikimate pathway. No feedback regulation by either Phe or Tyr was observed for either *AfCMCDT or *JbCDTCM (Table 4 and Figure S11).

**Table 4.**
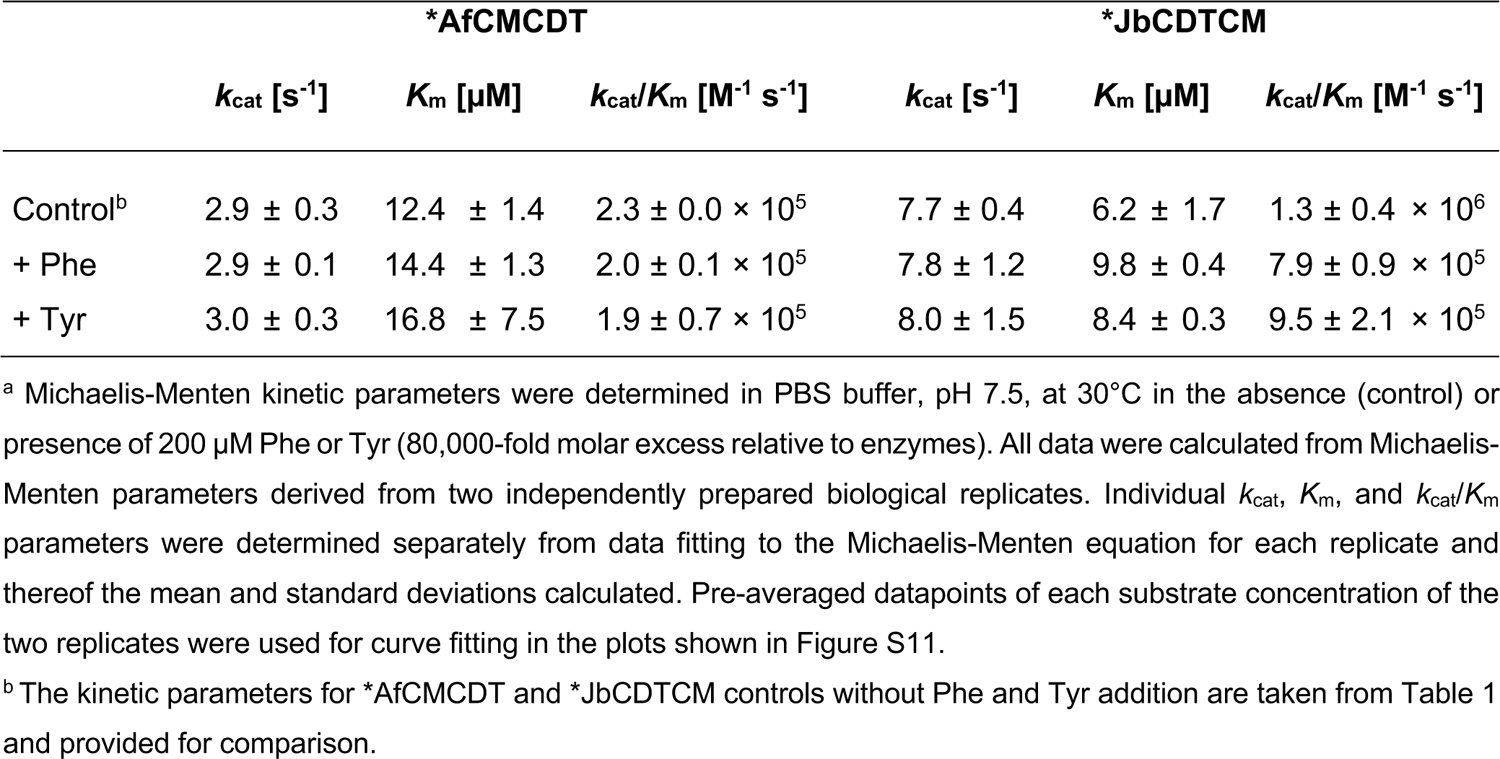
Test for feedback regulation of *AfCMCT and *JbCDTCM activity by Phe and Tyr^a^

|  | *AfCMCDT |  |  | *JbCDTCM |  |  |
| --- | --- | --- | --- | --- | --- | --- |
| | $k_{cat}$ [$s^{-1}$] | $K_m$ [$\mu M$] | $k_{cat}/K_m$ [$M^{-1} s^{-1}$] | $k_{cat}$ [$s^{-1}$] | $K_m$ [$\mu M$] | $k_{cat}/K_m$ [$M^{-1} s^{-1}$] |
| Control<sup>b</sup> | 2.9 ± 0.3 | 12.4 ± 1.4 | 2.3 ± 0.0 $\times 10^5$ | 7.7 ± 0.4 | 6.2 ± 1.7 | 1.3 ± 0.4 $\times 10^6$ |
| + Phe | 2.9 ± 0.1 | 14.4 ± 1.3 | 2.0 ± 0.1 $\times 10^5$ | 7.8 ± 1.2 | 9.8 ± 0.4 | 7.9 ± 0.9 $\times 10^5$ |
| + Tyr | 3.0 ± 0.3 | 16.8 ± 7.5 | 1.9 ± 0.7 $\times 10^5$ | 8.0 ± 1.5 | 8.4 ± 0.3 | 9.5 ± 2.1 $\times 10^5$ |

<sup>a</sup> Michaelis-Menten kinetic parameters were determined in PBS buffer, pH 7.5, at 30°C in the absence (control) or presence of 200 $\mu M$ Phe or Tyr (80,000-fold molar excess relative to enzymes). All data were calculated from Michaelis-Menten parameters derived from two independently prepared biological replicates. Individual $k_{cat}$ , $K_m$ , and $k_{cat}/K_m$ parameters were determined separately from data fitting to the Michaelis-Menten equation for each replicate and thereof the mean and standard deviations calculated. Pre-averaged datapoints of each substrate concentration of the two replicates were used for curve fitting in the plots shown in Figure S11.
<sup>b</sup> The kinetic parameters for \*AfCMCDT and \*JbCDTCM controls without Phe and Tyr addition are taken from Table 1 and provided for comparison.

### Assessing the genomic neighborhood of bifunctional fusion enzyme genes

To gain a better understanding of the potential biological role of these exported bifunctional enzymes, their genetic context was analyzed by using the webtool RODEO.^64^ RODEO stands for ‘Rapid Open reading frame Description and Evaluation Online’ and uses the NCBI accession number of the gene of interest to compile its surrounding 8 upstream and 8 downstream genes in the genomic neighborhood. Gene clusters in form of operons and closely aligned genes, particularly if they share the same orientation as those for the bifunctional fusion enzymes, may shed light on the enzymes’ biological roles in the organisms.

The genomic neighborhoods of all nine exported bifunctional fusion enzymes were analyzed using RODEO (Figure S12). In case of *AfCMCDT, four genes were allocated upstream with the same orientation of which the nearest two encode for a putative peroxidase and a hydrolase. The gene for *JbCDTCM is directly flanked by two genes in the same orientation to form an operon, where the gene upstream encodes a putative pyruvate carboxylase (HMGL-like) and downstream a putative lumazine-binding domain. These findings hint at potential roles of the bifunctional fusion enzymes in processes involving substrate degradation and binding. In the neighborhood of the genes for the other exported bifunctional fusion enzymes, an ABC transporter and a TonB receptor were found for *ScCMCDT, suggesting involvement in substrate uptake. An ATPase and another solute-binding protein are encoded directly downstream of the genes for *TaCMCDT and *TvCMCDT, whereas for *SbCMCDT and *SpCMCDT genes, no flanking coding regions with the same orientation were found. Interestingly, an intramembrane protease and an AI-2E family transporter are encoded by the two genes in the same direction and directly upstream of the one for *DsCDTCM, which suggests a role in signal recognition. Directly upstream of the *MpCDTCM gene is another substrate-binding protein encoded with the same orientation. In summary, we noticed a significant proportion of genes for substrate degradation and, particularly, for uptake and transport around the genes for the exported bifunctional fusion enzymes. Together with the discovery of an AI-2E transporter gene in the *DsCDTCM genomic neighborhood, encoding a transport protein for autoinducer 2 (AI-2)-type molecules, which are responsible for signaling and interspecies communication,^65, 66^ our analysis suggests that these exported bifunctional fusion enzymes may play a role in quorum sensing or interbacterial interactions.

## Discussion

Nine novel, very rare exported bifunctional fusion proteins consisting of an AroQ_γ_ CM domain and a CDT domain were discovered in a few γ- and β-proteobacterial species. Two distinct topologies were observed that divide these fusion enzymes into two distinct classes. In γ-proteobacteria, the CM domain is fused N-terminally to the CDT domain (*CMCDT), whereas in β-proteobacteria, the order is inverse (*CDTCM).

Seven of the nine exported bifunctional fusion enzymes were further investigated, and all showed CM and CDT activities in *E. coli* and in kinetic assays *in vitro*, except for *TaCMCDT, which showed no CDT activity at all in an *in vivo* complementation assay. Since *TaCMCDT could not be solubly overproduced in *E. coli* KA29 cells in spite of great efforts taken (data not shown), no reliable *in vitro* data could be generated for this enzyme. The low solubility is most likely also the explanation for the poor *in vivo* complementation (Figure S3). Production of the single split-off CDT domains yielded very little soluble protein for *AfCDT and none for *JbCDT, suggesting that the CDT domain benefits from the fused CM domain for stability in solution. The exposed interface to the partnering CM domain, possibly around the linker sequence, is probably the culprit for the low solubility after splitting the protein, given that *PaCDT with 40-60% sequence identity to the CDT domains was well produced in soluble form. This is further supported by the much more intricate domain interface found in *JbCDTCM (Figure 8C) than in *AfCMCDT (Figure 8B), explaining why it was even more difficult to produce and purify *JbCDT compared to *AfCDT.

To gain insight into the catalytic peculiarities of the fused CM and CDT domains, *in vitro* kinetic assays were performed. Michaelis-Menten parameters showed that *CDTCM variants exhibited slightly higher catalytic efficiencies than the *CMCDTs for the CM reaction (Table 1), consistent with their somewhat better growth during the *in vivo* complementation (Figure S3). More intriguing was the revelation that all tested bifunctional fusion enzymes showed a very low *K*_m_ (<7 µM) for CM activity (Table 1), which is at least 25-fold lower than the *K*_m_ of *MtCM (180 µM).^19^ Even though the catalytic rate constants for the CM domains are 2-20x lower compared to *MtCM (*k*_cat_ = 50 s^-1^), the catalytic efficiencies of the bifunctional fusion enzymes (*k*_cat_/*K*_m_ ≈ 1-3 × 10^6^ M^-1^ s^-1^) are still higher than that of *MtCM (*k*_cat_/*K*_m_ ≈3 × 10^5^ M^-1^ s^-1^). A very low *K*_m_ enables efficient binding of very low concentrations of chorismate, a property that would be a requirement for detection of chorismate if this molecule served as a signal, *e.g.*, for chemotaxis or inter-cellular communication. In comparison, the kinetic parameters for CDT activity are in the same order of magnitude and show similar catalytic efficiencies as *PaCDT (*k*_cat_/*K*_m_ ≈ 1 × 10^6^ M^-1^ s^-1^).^32^ This is consistent with the high degree of sequence identity and the conservation of all nine *PaCDT active site residues^32^ in the studied CDT domains.

The potential kinetic effects of the CM and CDT domain fusions were further addressed with coupled CM+CDT kinetic assays. While no catalytic benefit by the fusion was observed, it was noted that the CM domain’s *k*_cat_ was rate-limiting for the *CMCDT variants. For the *CDTCM enzymes, the *k*_cat_ for the combined CM+CDT reaction was slower than for either of the single-step CM or CDT catalytic reactions, suggesting that another process was rate-limiting. This could be a slow transition involving the steps from prephenate release by the CM domain to rebinding in the active site of the CDT domain, possibly due to topological or dynamic constraints in the fusion enzymes, which were shown to have diverging ligand exit/entry trajectories (Figure 6).

Additional insights were obtained from examining active-site KO and split single domain variants, using *AfCMCDT and *JbCDTCM as representatives for the *CMCDT and *CDTCM topologies, respectively. While CM activity was found to be unaffected compared to the wild-type fusion enzyme, a two-fold lower *K*_m_ for CDT activity was observed for *JbCDT and *AfCDT compared to their wild-type counterparts. This hints at competition between the CM and the CDT active sites for prephenate binding at low substrate concentration. The fact that mixing *AfCM and *AfCDT active site KO variants exhibited identical activity for the two-step process from chorismate to phenylpyruvate as the parental *AfCMCDT is a strong indication that having the domains covalently tethered brings no functional advantage. Whereas *MtCM and *PaCDT are known to form oligomers,^9,19, 32^ we show here by SEC that the bifunctional fusion enzymes are predominantly monomeric. This observation is supported by the protein crystal structures of *AfCMCDT and *JbCDTCM. Moreover, the crystal structures of all three bifunctional fusion enzymes presented here give no indication of substrate channeling. Taken together, there does not appear to be any catalytic benefit of the fused domains in these bifunctional enzymes, which raises the question what function these fusion enzymes have?

Domain fusions can have other benefits, for instance, it will facilitate coordinated gene expression or protein degradation.^67, 68^ Regulatory mechanisms to ensure stoichiometric concentrations of the active sites of the two sequential CM+CDT reactions must be put in place only once for the fusion protein, rather than for each enzyme separately. Also, both functions co-localize to the periplasm and the transfer across the membrane can occur simultaneously for CM and CDT. Even though not apparent in our kinetic assay, having co-localized active sites may be an advantage to sequester intermediates (here prephenate) from unwanted conversion or side reactions by competing enzymes.

It is still an enigma why these coupled enzymes are exported from the cytoplasm into a compartment where there is neither an obvious metabolic source for the substrate chorismate nor use of the products prephenate and phenylpuruvate. In fact, a role of these bifunctional fusion enzymes in the biosynthesis of aromatic amino acids is extremely unlikely; for one, they are exported to the periplasmic space and thus distant from the biosynthetic shikimate pathway.^1,2^ Moreover, they are not subjected to feedback regulation by the metabolic end products Phe or Tyr, which is otherwise ubiquitous in intracellular shikimate pathway enzymes.^1–3, 10, 16–18, 57–60^ The particularly low *K*_m_ values for the CM domains, which allow efficient binding of very low concentrations of the substrate chorismate, may hint at their true biological task. The genomic localization together with genes for solute-binding and transport proteins, as well as a presumed AI-2 transporter, suggests the involvement of the bifunctional fusion enzymes in processes such as chemotaxis, quorum sensing or interspecies communication.^65, 69, 70^ We postulate that the bifunctional fusion enzymes characterized here may be part of such systems.

## Materials and Methods

### Materials and general procedures

Chorismate was produced by a published procedure.^71^ All other chemicals were purchased from Sigma-Aldrich/Fluka (St. Louis, MO, USA). DNA manipulations were performed by standard procedures^72^ or Golden Gate Assembly (GGA) cloning (details in the Supplementary Materials and Methods). All DNA polymerases, DNA ladders, restriction endonucleases, and ligases were purchased from New England Biolabs (NEB; Ipswich, MA, USA). Sanger DNA sequencing and oligonucleotide synthesis were performed by Microsynth (Balgach, SG, Switzerland).

### Search for exported bifunctional fusion enzymes

The exported bifunctional fusion enzymes consisting of an AroQ_γ_ subclass CM domain and a cyclohexadienyl dehydratase domain were discovered by BLASTP^73^ using the fully translated open reading frame Rv1885c (*MtCM; UniProt: P9WIB9)^19^ including the signal peptide. The bifunctional fusion enzymes were identified by ∼50% query coverage but high sequence similarity and identity scores. Only search hits with full taxonomic classification to the species level were considered. The initially identified fusion enzymes were then themselves used for further BLASTP searches to find all nine final bifunctional fusion enzymes (for protein accession numbers, see Figure S12).

The amino acid sequences of the bifunctional fusion enzymes were reverse translated and codon-optimized for expression in *E. coli* using CLC Genomic Workbench v9.01 (QIAGEN^®^ CLC bio, Aarhus, Denmark). Furthermore, undesired restriction sites were removed manually by introducing silent mutations, and flanking restriction sites were introduced for cloning. A list of all nucleotide sequences can be found in the Supplementary Information (Figure S2). Gene synthesis was performed by GenScript (Piscataway, NJ, USA) or TWIST Bioscience (South San Francisco, CA, USA).

### Bacterial strains and plasmids

*E. coli* strains KA12^47, 48^ and KA29^19^ and plasmids pKTCTET-0^46^ and pKIMP-UAUC^48^ were previously described. The construction of pKTCTET derivatives encoding the bifunctional fusion enzymes with and without their natural signal sequences as well as the single domain and active site knockout variants of *AfCMCDT and *JbCDTCM are detailed in the Supplementary Materials and Methods.

### Protein overproduction and purification by Ni^2+^-NTA affinity chromatography

Luria Bertani (LB) starter cultures containing 100 µg/mL sodium ampicillin (Amp^100^) and 50 µg/mL kanamycin (Kan^50^) were prepared of KA29/pKTCTET derivatives for the appropriate cytoplasmatic bifunctional fusion enzyme variant and grown at 37°C, 230 rpm, overnight. The starter cultures were spun down and the pellet inoculated into fresh medium to remove β-lactamases in the supernatant before 800 mL LB production cultures containing 180 µg/mL sodium ampicillin (Amp^180^) and Kan^50^ were inoculated at a ratio of 1:100. The production cultures were incubated at 20°C, 160 rpm, for 2.5 days, then gently pelleted at 2,000 *g* for 10 min and resuspended in fresh 800 mL LB containing Amp^180^ and Kan^50^. The cultures were incubated for another 24 h at 20°C, 160 rpm. To harvest the cells the cultures were cooled on ice for 15 min and the cells pelleted by centrifugation at 4°C with 4,000 *g* for 15 min. The supernatant was discarded.

The pellets were resuspended with Sonication Buffer of 20 mM Tris-HCl, pH 8.0, containing 150 mM NaCl. Lysozyme was added to a final concentration of 1 mg/mL and the suspension incubated on ice for 30 min. The cells were lysed by sonication (3 rounds each with 100% amplitude, full cycle, 2 min sonication and 2 min breaks in an iced water bath, using an UP 200 s tip, Sonotrode S7, Dr. Hielscher GmbH, Teltow, Germany) and the lysates centrifuged at 4°C with 20,000 *g* for 20 min. The cleared supernatant was transferred to a beaker and Elution Buffer 2 (50 mM Tris-HCl, pH 8.0, containing 150 mM NaCl and 500 mM imidazole) was added to reach a final concentration of 20 mM imidazole. 6 mL of Ni^2+^-NTA slurry (Qiagen, Venlo, LI, Netherlands) were equilibrated in a gravity flow column with Wash Buffer (50 mM Tris-HCl, pH 8.0, containing 150 mM NaCl and 20 mM imidazole). The supernatant was loaded onto the metal affinity column and let flow through. The Ni^2+^-NTA bed was excessively washed with Wash Buffer before elution of metal-bound protein with 16 mL of Elution Buffer 1 (50 mM Tris-HCl, pH 8.0, containing 150 mM NaCl and 250 mM imidazole). The elution fractions were dialyzed using SnakeSkin tubing (Thermo Fisher Scientific, Waltham, MA, USA) with a molecular weight cut-off below the calculated mass of the protein. Dialysis was performed in 5 L Dialysis Buffer (20 mM Tris-HCl, pH 8.0, containing 150 mM NaCl) overnight at 4°C, followed by another dialysis in fresh 5 L Dialysis Buffer for 4-6 h at room temperature. The dialyzed protein was sterile filtered (0.22 µm), stored at 4°C for direct use or snap frozen in liquid nitrogen for storage at −80°C.

Enzyme concentrations [E] were determined using the Bradford assay.^74^ The expected molecular masses were confirmed by electrospray ionization mass spectrometry (ESI-MS) at MoBiAS (Laboratory of Organic Chemistry, ETH Zurich, Switzerland) with mass data reported in Table S2.

### Size-exclusion chromatography

Analytical size-exclusion chromatography (SEC) was performed using the BioLogic DuoFlow™ system with a QuadTec™ UV/Vis Detector (BioRad Laboratories Inc., Hercules, CA, USA) and the Superdex™ 200 increase 10/300 GL column (Cytiva, Little Chalfont, UK). 20 mM Tris-HCl, pH 8.0, containing 150 mM NaCl degassed and prechilled to 4°C was used to equilibrate the whole system and as running buffer. 0.8-1.3 mg of samples were injected in a volume of 0.6 mL at a flow rate of 0.5 mL/min for injection, and passed through the column with 25 mL running buffer at a flow rate of 0.7 mL/min. The monitored absorbance at 214 nm (A_214nm_) was used to plot elution profiles of the protein standard mix Supelco^®^ 69385 Protein Standard Mix 15-600 kDa (Sigma-Aldrich/Merck, Darmstadt, Germany), and the analyzed protein samples.

### CM, CDT, and coupled CM-CDT *in vitro* kinetic assays

For the final read-out, the absorbance of phenylpyruvate after conversion to its enolate form was recorded for each the CM, the CDT, and the CM-CDT coupled discontinuous assay, as detailed in Figure S13 and Table S3. The enolate form of phenylpyruvate exhibits a high extinction coefficient (ε) of 17,500 M^-1^ cm^-1^ at 320 nm, which allows the detection of very low phenylpyruvate concentrations. Six different substrate concentrations ranging from 2.5-100 µM, each at 4 time points between 0-4 min were measured. A more detailed experimental protocol can be found in the Supplementary Materials and Methods. From the four time points, the *v*_init_ value for each substrate concentration was calculated, with additional correction for the spontaneous background turnover rate at 30°C for chorismate or prephenate. The Michaelis-Menten graphs were generated using Prism (GraphPad Software, San Diego, CA, USA) by plotting *v*_init_/[E] against the substrate concentration [S]. Data points were fitted to the Michaelis-Menten equation *v*_init_ = *k*_cat_ × [E] × [S] / (*K*_m_ + [S]) to determine the catalytic rate constant *k*_cat_ and the Michaelis constant *K*_m_. All reported catalytic parameters are averages with standard deviations (σ_n-1_) calculated from individually determined values for *k*_cat_, *K*_m_, and *k*_cat_/*K*_m_ of Michaelis-Menten kinetics performed separately with two independently prepared biological replicates.

Feedback regulation by either Phe or Tyr was analyzed by performing the same protocol as for the discontinuous coupled CM+CDT assay, but with additional 200 µM Phe or 200 µM Tyr in the reaction buffer.

### Protein crystallography

#### Crystallization

*AfCMCDT crystallized at 20 °C in a sitting-drop 96-well SWISSCI UVXPO 2-lens crystallization plate. Diffraction quality crystals grew from a 1:3 volume ratio (150 nL:450 nL) mixture of 5 mg/mL protein, stored in 20 mM Tris-HCl, pH 8.0, 150 mM NaCl, and condition G7 from the PACT premier crystallization screening formulation (Molecular Dimensions Ltd.), containing 0.1 M Bis Tris propane, pH 7.5, 0.2 mM sodium acetate and 20% w/v PEG3350, at 20°C. The crystal was cryoprotected by quickly dipping it in a crystallization solution complemented with glycerol to a final concentration of 20% v/v glycerol, and was thereafter flash-cooled in liquid nitrogen and stored until data collection. 4 mg/mL (85 µM) *JbCDTCM in 20 mM Tris-HCl, pH 8.0, was used for crystallization experiments. *JbCDTCM crystals were obtained at 20 °C from a 1:1 (200 nL:200 nL) mixture of protein sample and solution C1 from the Morpheus crystallization screening kit (Molecular Dimensions Ltd.), containing 30% w/v PEG500MME_P20K (10% w/v PEG 20000, 20% v/v PEG 500 monomethyl ether), 0.09 M NPS (0.3 M sodium nitrate, 0.3 M disodium hydrogen phosphate, 0.3 M ammonium sulfate), and 0.1 M MES/imidazole, pH 6.5. The crystallization experiment was set up in a sitting-drop 96-well UVXPO 2-lens crystallization plate (SWISSCI). Crystals were flash-cooled in liquid nitrogen without using any additional cryoprotectant and stored for data collection.

*DsCDTCM crystals grew at 4 °C from a 1:1 (300 nL: 300 nL) mixture of 5.6 mg/mL protein in 20 mM Tris-HCl, pH 8.0, and condition G12 from the JCSG+ crystallization screening kit (Molecular Dimensions Ltd.), containing 3 M NaCl and 0.1 M Bis Tris, pH 5.5, in a sitting drop 96-well SWISSCI UVXPO 2-lens crystallization plate. The crystal was cryoprotected by quick soaking in freshly prepared reservoir solution complemented with ethylene glycol to a final concentration of 20% v/v ethylene glycol, flash-cooled in liquid nitrogen and stored for data collection.

#### Data collection, structure determination, and refinement

Data collection for *AfCMCDT was performed at 100 K at the European Synchrotron Radiation Facility (ESRF, Grenoble) beamline ID30A-1, which was equipped with a PILATUS3 2M detector. Diffraction data for *JbCDTCM and *DsCDTCM were collected at 100 K at the BioMAX beamline, MAX-IV (Lund, Sweden), equipped with an EIGER X 16M detector. Data sets covering 360° were collected for each crystal, with oscillation ranges of 0.2° for *AfCMCDT, and 0.1° for *JbCDTCM and *DsCDTCM. The resolution at the detector edge was set to 1.0 Å for *AfCMCDT, to 1.5 Å for *JbCDTCM, and to 1.8 Å for *DsCDTCM, respectively. All three data sets were indexed and integrated using automated pipelines at the synchrotrons (*autoPROC*^75^). All data sets showed some anisotropy, and were scaled and merged with the *STARANISO* server.^75^

The structures were solved by molecular replacement with *Phaser,*^76^ a software from the *CCP4* program suite.^77^ As search models, the CM from *Burkholderia thailandesis* (*BtCM; PDB ID: 6CNZ)^66^, and CDT from *P. aeruginosa* were used for both *AfCMCDT and *JbCDTCM. *BtCM shares 45% and 32% sequence identity with *JbCM and *AfCM, respectively, and *PaCDT shares 62% and 50% sequence identity with *JbCDT and *AfCDT, respectively. The *DsCDTCM structure was solved using the *JbCDTCM structure as search model.

The structures were refined by alternating cycles of maximum-likelihood refinement using *REFMAC*^78^ (a program from the *CCP4* software suite)^77^ with model inspection and rebuilding using *Coot*.^79^ Water molecules, alternative conformations, and ligands were added to the structures in later refinement cycles, interpreting peaks in the σ_A_-weighted m*Fo-*D*Fc* difference electron density map, taking the content of the mother liquor into account. Sodium ions were added when the difference density peaks were in close proximity (closer than a typical H-bond) to one or several carbonyl groups, or if an octahedral coordination was observed (one instance). Chloride ions were modeled when placement of a water molecule led to artificially low *B*-factors and residual difference electron density at the exact position (and if the chemical environment was compatible with a negative charge). Some of the chloride ions were found in the CDT active site, at a position likely binding the carboxylate groups of the substrate (prephenate) or product (phenylpyruvate; Figure 7C). In the CDT active site of *JbCDTCM, positive difference density revealed the presence of a MES molecule from the crystallization buffer; the sulfonate group occupied the position of chloride ions in *DsCDTCM. Acetate, also a component of the crystallization solution, was modeled into both CM and CDT active sites of *AfCMCDT. Finally, occupancy refinement was carried out with *phenix.refine*,^80^ a tool from the *PHENIX* software package.^81^ Data collection, processing and refinement statistics (as calculated with *PHENIX*^81^) are summarized in Table 2 and Table S1.

## Accession Numbers

The atomic coordinates and structure factors of the crystal structures of *AfCMCDT (PDB ID: 8CQ3) *JbCDTCM (PDB ID: 8CQ4), and *DsCDTCM (PDB ID: 8CQ6) have been deposited in the Protein Data Bank.

## Appendix A. Supplementary Data

Supplementary Information to this article can be found online.

## Acknowledgements

We would like to thank Michele Cascella for supervising the simulations. X-ray diffraction experiments were performed on beamline ID30A-1 at the European Synchrotron Radiation Facility (ESRF), Grenoble, France and on the BioMAX beamline at MAX IV, Lund, Sweden. We are grateful to Local Contacts Didier Nurizzo and Oskar Aurelius, Johan Unge at the ESRF and at MAX IV, respectively, for providing assistance in using the beamlines. This work was funded by a grant from the Swiss National Science Foundation to P.K. (grant 310030M_182648). We acknowledge services provided by the MoBiAS facility, Laboratory of Organic Chemistry, ETH Zurich, and use of the UiO Structural Biology Core Facilities, which are part of the Norwegian Infrastructure NORCRYST, supported by the Norwegian Research Council (grant 245828).

## Author Contributions

P.K. conceived the study. C.S., T.K., K.W.R., G.C. and U.K. were additionally involved in the planning of the experiments. C.S. and K.W.R. performed the bioinformatics analyses, designed and assembled the recombinant constructs. C.S. established protein production and purification conditions, carried out the complementation tests, engineered and purified all variants, and performed the kinetic analyses and the SEC investigation. C.S. and K.W.R. were supervised by P.K.. T.K. crystallized the proteins and solved and refined the X-ray structures, supervised by G.C. and U.K., who also validated the crystal structures. C.S. wrote the first draft of the manuscript, and T.K. contributed the first version of the crystallographic analysis. All authors contributed to the revision of the manuscript.

## Abbreviations Footnote

(*)CDT: (exported) cyclohexadienyl dehydratase

(*)CM: (exported) chorismate mutase

*AfCDTCM: *CDTCM of *Aequoribacter fuscus*

*AfCM and *AfCDT: split domain variants of *AfCDTCM

*CDTCM: exported bifunctional fusion enzyme with CDT domain N-terminal to CM domain

*CMCDT: exported bifunctional fusion enzyme with CM domain N-terminal to CDT domain

*DsCDTCM: *CDTCM of *Duganella sacchari*

*JbCDT and *JbCM: split domain variants of *JbCDTCM

*JbCDTCM: *CDTCM of *Janthinobacterium* sp. HH01

*MpCDTCM: *CDTCM of *Massilia phosphatilytica*

*MtCM: exported CM of *Mycobacterium tuberculosis* (AroQγ subclass)

*PaCDT: exported CDT of *Pseudomonas aeruginosa* (also called PheC)

*SbCMCDT: *CMCDT of *Shewanella baltica*

*ScCMCDT: *CMCDT of *Steroidobacter cummioxidans*

*TaCMCDT: *CMCDT of *Thalassomonas actiniarum*

a.u.: asymmetric unit

CM+CDT: coupled CM and CDT kinetic activity measurement

CM and CM: fusion enzyme variants with knocked-out CDT activity

CDT and CDT: fusion enzyme variants with knocked-out CM activity

DAHP: 3-desoxy-_D_-*arabino*-heptulosanate 7-phosphate

EcCM: intracellular CM of *Escherichia coli* (AroQα subclass; also called P-protein)

KO: knocked-out catalytic active site

MES: 2-(*N*-morpholino)ethanesulfonic acid

PDB: Protein Data Bank

PDH: prephenate dehydrogenase

PDT: Prephenate dehydrogenase

r.m.s.d.: root mean square difference/deviation

RODEO: Rapid Open reading frame Description and Evaluation Online

TSA: *endo*-oxabicyclic transition state analog of the CM reaction

