## Supplementary Information for "Novel exported bifunctional fusion enzymes with chorismate mutase and cyclohexadienyl dehydratase activity: shikimate pathway enzymes teamed up in no man’s land"

### Supplementary Materials and Methods

#### Construction of pKIMP-UA

The helper plasmid pKIMP-UA was generated from the plasmid pKIMP-UAUC by restriction digestion of with *Sal*I and *Xho*I that flank the promoter and the *pheC* gene. The digested fragments were separated on an agarose gel and the larger 3733 bp fragment with the vector backbone was cut out and the DNA extracted using the Zymoclean™ Gel DNA Recovery kit (Zymo Research Corporation, Irvine, CA, USA). Because *Sal*I and *Xho*I restriction digestions result in compatible overhangs the open vector fragment could be circularized to form the fully functional plasmid pKIMP-UA (3733 bp). The ligation mix was transformed into SEM[1] prepared chemically competent KA12[2] cells and the mixture plated onto LB agar plates containing 30 µg/mL chloramphenicol ($Cam^{30}$). Successful cloning was checked by a single site restriction digestion analysis on an agarose gel for the correct size.

#### *In vivo* complementation assay

The pKTCTET plasmids for the expression of the bifunctional enzymes genes without their signal sequence were transformed into SEM chemically competent KA12/pKIMP-UAUC[3] cells for an *in vivo* assay for complementation of CM activity or KA12/pKIMP-UA cells for an *in vivo* coupled CM+CDT complementation. Transformants were plated onto agar plates with addition of 100 µg/mL sodium ampicillin ($Amp^{100}$) and $Cam^{30}$. A purification streak-out onto a fresh LB agar plate containing $Amp^{100}$ and $Cam^{30}$ was performed, and single colonies were used for the complementation assays. They were streaked out onto M9c minimal medium[4] agar plates containing $Amp^{100}$ and $Cam^{30}$ with addition of either 20 µg/mL L-Phe and 20 µg/mL L-Tyr (+FY), 200 or 50 ng/mL tetracycline ($Tet^{200}$ or $Tet^{50}$, respectively), or none. The plates were incubated at 30°C for up to 5 days after wrapping them in saran foil to prevent drying of agar. Every day the plates were analyzed and the growth of colonies rated on an arbitrary scale from 0 to 9 [0, no trace of growth; 1, some cell material at start of streak out; 2, cell material along the first streak out lane; 3, single colonies visible as dots against the light; 4, tiny single colonies (ø < 0.5 mm); 5, small single colonies (ø ≈ 0.5 mm); 6, small single colonies that are easy to pick (ø ≈ 1 mm); 7, middle sized single colonies (ø ≈ 2 mm); 8, large single colonies (ø ≈ 3 mm); 9, giant single colonies (ø > 3 mm)].

Cloning of the bifunctional fusion enzymes and variants into vector pKTCTET

The protein sequences of *AfCMCDT, *SbCMCDT, and *TaCMCDT (accession numbers in Figure S12) were identified as detailed in the manuscript. The sequences were reverse translated and codon optimized for *Escherichia coli* by CLC Genomics Workbench v9.01 (QIAGEN® CLC bio, Aarhus, Denmark). Silent restriction sites were introduced after the signal sequence and between the CM and CDT domains. The genes flanked by tandem *Eco*RI/*Nde*I and *Xho*I/*Hin*dIII sites were ordered from GenScript (Piscataway, NJ, USA) cloned in a pUC vector backbone in the *Eco*RI and *Hin*dIII restriction sites. The synthesized genes were cut out from the pUC vectors employing *Nde*I and *Xho*I restriction enzymes, yielding 1251 bp, 1266 bp, and 1290 bp fragments for *AfCMCDT, *SbCMCDT, and *TaCMCDT genes, respectively. The genes were individually ligated to the correspondingly cut pKTCTET-0[4] vector (2797 bp) yielding 4048 bp (pKTCTET-AfCMCDT-HC), 4063 bp (pKTCTET-ScCMCDT-HC), and 4087 bp (pKTCTET-TaCMCDT-HC) plasmids. The plasmids encoding the respective *CMCDT variants including the native secretion tag and a C-terminally fused $His_6$-tag stemming from vector pKTCTET-0 that is appended *via* a Leu-Glu linker (CTCGAG, *Xho*I restriction site) for metal affinity chromatography (the sequences of the *Nde*I/*Xho*I-flanked genes are given in Figure S2 below).

The genes for the other exported bifunctional fusion enzymes were designed from the amino acid sequence (accession numbers in Figure S12) by reverse translation and codon optimization for production in *E. coli* using CLC Genomics Workbench v9.01. Gene synthesis was done at TWIST Bioscience (South San Francisco, CA, USA) and the genes (Figure S2) were cut out from the received TWIST carrier plasmids using *Nde*I and *Xho*I restriction digestion for ligation into the correspondingly cut 2797 bp acceptor fragment of vector pKTCTET-0 as described above.

All ligation products were transformed into SEM chemically competent BL21-Gold(DE3) or KA12 cells and the constructs verified by DNA sequence analysis to confirm the gene sequences of plasmids pKTCTET-AfCMCDT-HC (4048 bp), pKTCTET-ScCMCDT-HC (4081 bp), pKTCTET-TaCMCDT-HC (4087 bp), pKTCTET-SbCMCDT-HCT (4063 bp), pKTCTET-JbCDTCM-HC (4051 bp), pKTCTET-DsCDTCM-HC (4024 bp), and pKTCTET-MpCDTCM-HC (4063 bp).

### Construction of pKTCTET plasmids encoding the cytoplasmic bifunctional enzyme variants

The pKTCTET plasmids encoding the exported bifunctional enzyme constructs were taken as DNA templates to amplify the genes for the bifunctional enzymes without their signal peptide by PCR. The primer pairs used were #0673 AfCM_f (5'-AATCAT-ATGGACAATCATTCGGAGCAGACG) and #0706 pKTCTET-0_ext_r (5'-GGAA-TAAGGGCGACACGG) (1544 bp fragment size) for pKTCTET-AfCMCDT-HCT (3985 bp), #0828 ScCM_f (5'-TATCATATGGCGTCATTTACAGGTCCCAAC) and #0706 (1577 bp fragment size) for pKTCTET-ScCMCDT-HCT (4018 bp), #0672 TaCM_f (5'-AATCAT-ATGCAGACCCAAGCCAATGAACC) and #0706 (1547 bp fragment size) for pKTCTET-TaCMCDT-HCT (3988 bp), #0671 SbCM_f (5'-AATC-ATATGGATACGGACGAAAACGCG-GAG) and #0706 (1532 bp fragment size) for pKTCTET-SbCMCDT-HCT (3973 bp), #0826 JbCDT_f (5'-TATCATATGGATCATAGGCTGGATGACATCAC) and #0706 (1562 bp fragment size) for pKTCTET-JbCDTCM-HCT (3967 bp), #0825 DsCDT_f (5'- TATCAT-ATGGGCCGTTTGGAAGAGATTC) and #0706 (1529 bp fragment size) for pKTCTET-DsCDTCM-HCT 3970 bp), as well as #0827 MpCDT_f (5'-TATCATATGGGTCATTTA-GATGATATTGCCGC) and #0706 (1550 bp fragment size) for pKTCTET-MpCDTCM-HCT (3991 bp). With the Zymo DNA Clean & Concentrator-5 kit (Zymo Research Corporation, Irvine, CA, USA) the amplified PCR products were purified and digested with *Nde*I and *Xho*I. This removed a 6 bp fragment upstream and a 350 bp fragment downstream of the gene, which were eliminated by purification over an agarose gel. The digested PCR fragments were cut out and the DNA was extracted using the Zymoclean™ Gel DNA Recovery kit. The digested fragments were ligated with the *Nde*I and *Xho*I cut (2797 bp) pKTCTET-0 acceptor fragment and transformed into SEM chemically competent KA12 cells. The constructs were verified by DNA sequence analysis using the T7 primer.

### Construction of pKTCTET-sfGFP (GGA)

For facilitated cloning of further variants of *AfCMCDT and *JbCDTCM, Golden Gate Assembly (GGA)[5] compatible pKTCTET-0 plasmid variant (pKTCTET-sfGFP with 3509 bp size and a gene encoding the superfolder green fluorescent protein (sfGFP) as insert) was constructed. Therefore, two undesired *Bsa*I restriction sites, one located in the T7 promoter sequence and one in the ampicillin resistance gene, had to be removed. Three PCRs using pKTCTET-0 as template were performed with the primer pairs: #0993 pKTCTET-1_f (5'-

TGGCCTTTTGCTCACATGCTTAAGACCC) and #0994 pKTCTET-1_r (5'-ATTTCTA-GAGGGAAACCGTTGTGGTCACCC) (787 bp fragment size), #0995 pKTCTET-2_f (5'-TAAGGTCTCCCCGCGGTATCATTGCAGCACTGGG) and #0996 pKTCTET-2_r (5'-TGGGTCTTAAGCATGTGAGCAAAAGGCC) (982 bp fragment size), as well as #0997 pKTCTET-3_f (5'-CTGCCACCGCTGAGCAATAACTAGC) and #0998 pKTCTET-3_r (5'-TAAGGTCTCCGCGGGACCCACGCTCACC) (956 bp fragment size). The PCR products were cleaned by the DNA Clean and Concentrator-5 kit and each digested with a different set of two restriction enzymes. *Xba*I and *Afl*II for #0993-#0994 (desired fragment with 760 bp size), *Afl*II and *Bsa*I for #0995-#0996 (desired fragment with 959 bp size), and *Bsa*I and *Blp*I for #0997-#0998 (desired fragment with 933 bp size). The desired fragments were purified again using the DNA Clean and Concentrator-5 kit. The acceptor vector pKTCTET-0 was restriction digested with *Xba*I and *Blp*I (desired fragment with 1432 bp size) and purified by agarose gel electrophoresis and the DNA extracted using the Zymoclean™ Gel DNA Recovery kit. All fragments were ligated together using standard T4 DNA ligase and then transformed into SEM chemically competent KA12 cells. The sequence of the generated intermediate plasmid without *Bsa*I restriction sites, but still without the sfGFP gene, was checked by DNA sequencing with the external primers covering the entire PCR'ed region.

The intermediate plasmid was also digested with *Xba*I and *Blp*I to generate the 2664 bp acceptor vector. Both fragments were purified *via* agarose gel electrophoresis, extracted using the Zymoclean™ Gel DNA Recovery kit and ligated with T4 DNA ligase. The ligation product was transformed into SEM chemically competent KA29[6] cells and successful cloning confirmed by sfGFP gene expression.

The gene for sfGFP was taken from plasmid pKINT-sfGFP by restriction digestion with *Xba*I and *Blp*I to generate the 845 bp sfGFP fragment. pKINT-sfGFP had been created by amplifying the gene for the C-terminally His-tagged superfolder GFP (sfGFP) from template pET29b(+)_sfGFP-6His (kindly provided by Moritz Pott and similar to pET29b(+)_sfGFP(133AMBER)-6His, except for the PCR assembly using oligonucleotides o28 and o29),[7] using the primer pair #0991 sfGFP_BsaI_f (5'-TTTCATATGGAGACCAG-CAAAGGAGAAGAACTTTTCACTGGAGTTG) and #0992 sfGFP_BsaI_r (5'-TTTAAGCTT-GAGACCTTAGTGGTGGTGGTGGTGGTGCTC). The PCR-amplified 774 bp fragment incorporated *Nde*I/*Bsa*I and *Bsa*I/*Hin*dIII restriction sites flanking the sfGFP gene. This PCR product and pKINT-0[8] as acceptor vector were each digested with *Nde*I and *Hin*dIII and then purified over an agarose gel. The 762 bp *Nde*I/*Hin*dIII-digested sfGFP gene and the 3478 bp acceptor fragment were isolated using the Zymoclean™ Gel DNA Recovery kit and ligated using T4 DNA ligase. The ligation was transformed into SEM chemically competent KA12 cells and plated onto a LB agar plate containing 50 µg/mL kanamycin. Successful assembly of pKINT-sfGFP (4240 bp) was confirmed by DNA sequencing with the T7 primer.

### Golden Gate Assembly (GGA) cloning protocol

Oligonucleotides for PCR were designed to include the *Bsa*I restriction site (5'-GGTCTC) on both ends to generate the desired overhang. The PCR products were directly cleaned up using the DNA Clean and Concentrator-5 kit and then restriction digested with *Dpn*I performed to eliminate the template plasmid DNA used for the PCR. The restriction digestions were again cleaned up using the Zymo DNA Clean and Concentrator-5 kit, and the DNA concentration was determined using a NanoDrop device (Waltham, MA, USA).

A GGA reaction mixture was composed of 20-50 ng of digested and purified PCR products, 20-50 ng of the GGA-compatible plasmid pKTCTET-sfGFP, 0.5 µL BsaI-HF®v2 (#R3733), 0.5 µL T7 DNA ligase (#M0318), and d$H_2O$ to a final volume of 20 µL. The GGA reaction cycles encompassed incubation steps of 60 s at 37°C followed by 60 s at 16°C for a total of 30 cycles, a last step consisted of a 10 min incubation at 55°C and then cooling to 4°C. The entire 20 µL of GGA reaction was transformed into 200 µL of SEM chemically competent cells. Any unsuccessful GGA cloning can easily be recognized because it would lead to production of the visually detectable sfGFP protein.

### Construction of active site KO and single, split-off domain variants of *AfCMCDT

The plasmid pKTCTET-AfCMCDT-HCT was used as DNA template for the PCRs. The CM active site KO variant pKTCTET-AfCMCDT-HCT_K48A (3963 bp) was assembled with the primer pairs #1034 AfCM_f2 and #1035 MgCMCDT_K48A_r (5'-TGGTCTCCATGCG-TATTTT¬GCCACTTCG-GGCATCAATG) (100 bp fragment size) as well as #1036 AfCMCDT_K48A_f (5'-TGGTCTCCGCATGGCATCACAACCTGCCAATCG) and #1039 AfCDT_r (5'-TGGTCTCAAGCTTATTAGTGGTGGTGGTGGTGGTGC) (1156 bp fragment size). For the CDT active site KO variant pKTCTET-AfCMCDT-HCT_E353Q (3963 bp) the primer pairs #1034 and #1037 AfCMCDT_E353Q_r (5'-TGGTCTCGCTTGTATGC-TATCGGTGAACATGGCATCGG) (1015 bp fragment size) as well as #1038 AfCMCDT_E353Q_f (5'-TGGTCTCACAAGCCCAGTTGCAAGCGACAAAAC) and #1039 (241 bp fragment size) were used. The PCR products were directly cleaned up using the Zymo

DNA Clean and Concentrator-5 kit, followed by restriction digestion with *Dpn*I of the PCR template. After another purification using the DNA Clean and Concentrator-5 kit, the two PCR products of the AfCMCDT-HCT_K48A cloning or the AfCMCDT-HCT_E353Q cloning were assembled into pKTCTET-sfGFP *via* GGA described above.

The split-off CM domain variant pKTCTET-AfCM-HNT (3277 bp) was assembled with the primer pair #0678 His6-AfCM_f (5'-AATCATATGCACCATCATCATCATCATTCTTCTGGTG-ACAATCATTCGGAGCAGAC) and #0704 AfCM_r (5'-AATCTACTAGTCATTATTACAAGTT-GGGCGGGGTAAG) (526 bp fragment size), and pKTCTET-AfCDT-HCT (3514 bp) with #0703 AfCDT_f (5'-ATGTACATATGGATCCTAGGGATACTCTCGC) and #0706 pKTCTET-ext_r (5'- GGAATAAGGGCGACACGG) (1075 bp fragment size). The PCR products were directly purified using the Zymo DNA Clean & Concentrator-5 kit before restriction digestion with *Nde*I and *Spe*I, which cut off 6+8 bp flanking the AfCM-HNT and 8+318 bp flanking the AfCDT-HCT PCR products. The acceptor vector pKTCTET-0 was restriction digested with *Nde*I and *Spe*I to generate the cut 2765 bp pKTCTET acceptor fragment. All digestion reaction products were purified *via* agarose gel electrophoresis, the desired band cut out and the DNA extracted using the Zymoclean™ Gel DNA Recovery kit. The cut acceptor vector and the according cut PCR products for AfCM-HNT or AfCDT-HCT were ligated using the T4 DNA ligase.

All ligations were transformed into SEM chemically competent KA29 cells and plated onto LB agar plates containing $Amp^{100}$ and $Kan^{50}$. The constructs were controlled by DNA sequence analysis using the T7 primer.

### Construction of active site KO and split-domain variants of *JbCDTCM

The active site KO and split-domain variants were assembled by PCR using pKTCTET-JbCDTCM-HCT as template DNA. For the CDT active site KO variant pKTCTET-JbCDTCM-HCT_E200Q (3967 bp) the primer pairs #0705 and #1001 JbCDTCM_E200Q_r (5'-TGGTCTCTCTGAATTGCATCCGTGATCATCAAATCTG) (783 bp fragment size) as well as #1000 JbCDTCM_E200Q_f (5'-TGGTCTCTTCAGACGCGGCTGCAACAAC) and #0706 (1016 bp fragment size) were used, and for the CM active site KO variant pKTCTET-JbCDTCM-HCT_K287A (3967 bp) the primer pairs #0705 and #1003 JbCDTCM_K287A_r (5'-TGGTCTCCACGCAGCACGGGCAACGGC) (1045 bp fragment size) as well as #1002 JbCDTCM_K287A_f (5'-TGGTCTCTGCGTGGAACGTGCAGGCTCC) and #0706 (754 bp fragment size) were used. The PCR products were purified using the DNA Clean &

Concentration-5 kit and the ones generated with the #0705 primer restriction digested with *Xba*I and *Bsa*I PCR products involving the #0705 primer, which cut off 220+10 bp flanking the fragments, whereas the PCR products generated with the #0706 primer *Bsa*I and *Spe*I were used, cutting off 10+318 bp flanking the fragments. pKTCTET-0 as acceptor vector was restriction digested with *Xba*I and *Spe*I. All restriction digested reactions were purified by agarose gel electrophoresis and the DNA extracted using the Zymoclean™ Gel DNA Recovery kit. The purified digested fragments for JbCDTCM-HCT_E200Q or JbCDTCM-HCT_K287A, were ligated simultaneously with the cut 2726 bp long pKTCTET acceptor fragment.

The split CM domain variant pKTCTET-JbCM-HCT was assembled with the primer pair #1006 JbCM_f (5'- TGGTCTCCATATGGGCTTGGAACCCTTACGTTTG) and #1007 JbCM_r (5'-TTTCTCTAAGCTTA-TTAGTGGTGGTGGTGGTGG) (513 bp fragment size), and the split CDT domain pKTCTET-JbCDT-HNT_WLDFPW with #1004 His6-JbCDT_f (5'-TGGTCTCCATATGCACCATCATCATCACCACGATCATAGGCTGGATGACATCACG) and #1005 JbCDT_r2 (5'-TGGTCTCAAGCTTTATTACCAGGGAAAATCCAGCCAC) (754 bp fragment size). The PCR products were purified using the DNA Clean & Concentrator-5 kit, and the PCR template DNA restriction digested using *Dpn*I. After another purification using the DNA Clean & Concentrator-5 kit and the two PCR products of the JbCM-HCT cloning or the JbCDT-HNT cloning were ligated into the plasmid pKTCTET-sfGFP *via* GGA.

All ligations were transformed into SEM chemically competent KA29 cells and plated onto LB agar plates with $Amp^{100}$ and $Kan^{50}$ agar plates. Successful cloning was confirmed by DNA sequencing with the T7 primer.

### Detailed CM, CDT, and coupled CM-CDT *in vitro* kinetic assay protocol

All three performed discontinuous assays have the same final readout of absorbance at 320 nm ($A_{320nm}$) of phenylpyruvate in its enolate form (Figure S13).

For the Michaelis-Menten-based kinetic analysis, six different substrate concentrations (2.5-100 µM), each with 4 different incubation periods (0-4 min) were measured resulting in a total of 24 individual reactions (Table S3). To keep the substrate turnover below 25%, the threshold assumed to still give initial velocities ($v_{init}$) needed for Michaelis-Menten kinetics, the incubation periods at the lower substrate concentrations were set shorter than 1 min. The $v_{init}$ values were calculated with the slope of the substrate consumption curve from the four time points for each substrate concentration. Furthermore, the calculated $v_{init}$ for each substrate concentration was corrected for the spontaneous background turnover rate at 30°C for chorismate ($1.15 \times 10^{-5}$ $s^{-1}$) or prephenate ($2.5 \times 10^{-5}$ $s^{-1}$) and then divided by the enzyme concentration to plot $v_{init}$/[E] against the substrate concentration. Michaelis-Menten curves were fitted through the data points and to calculate the rate constant $k_{cat}$ and the Michaelis constant $K_m$ using Prism (GraphPad Software, San Diego, CA, USA). All reported catalytic parameters are the average derived from the full kinetic analysis of two independently prepared biological replicates.

A single reaction volume was 200 µL containing 190 µL Reaction Buffer (50 mM potassium phosphate, pH 7.5, containing 0.1 mg/mL BSA and the corresponding chorismate or prephenate concentrations; Table S3) and 10 µL of the enzyme. For accurate reaction timings, 950 µL master reaction mixes for each substrate concentration, but without the addition of enzyme were pipetted, enough for 5 reactions to ensure identical substrate and enzyme concentrations for 4 reaction time points and some backup mixture to avoid pipetting errors. For the 0 min time point, the 190 µL of reaction mixture were pipetted from the master reaction mix into a microtube containing 100 µL of 2 M HCl (CM assay) or 200 µL of 5 M NaOH (CDT and CM+CDT assays). For all other time points, the 40 µL of the appropriate enzyme was added to the remaining 760 µL of master reaction mix. The reaction mix was quickly vortexed and put into a water bath at 30°C. At each time point, 200 µL of the reaction solution were transferred into a microtube containing 100 µL of 2 M HCl (CM assay) or 200 µL of 5 M NaOH (CDT and CM+CDT assays) and quickly vortexed to quench the reaction. For the CM assay the full chemical conversion of prephenate to phenylpyruvic acid was reached after 10 min incubation at 30°C, and then 100 µL of 10 M NaOH were added to establish an alkaline pH. After processing of all 24 reactions, the 10 µL of the appropriate enzyme was added to the 0 min time point reactions to ensure identical assay compositions for all absorbance measurements. All reactions in the microtubes were centrifuged for 1 min at room temperature at 20,000 *g* and the absorbance was measured at 320 nm ($A_{320nm}$) in 0.5 mm quartz cuvettes.

### Supplementary Figures

**(A)**

| Identity \ Similarity | MtCM | AfCM | ScCM | TaCM | TvCM | SbCM | SpCM | JbCM | DsCM | MpCM |
|---|---|---|---|---|---|---|---|---|---|---|
| MtCM | | 18.29 | 24 | 17.68 | 16.29 | 21.02 | 17.61 | 24.86 | 23.33 | 23.2 |
| AfCM | 29.14 | | 22.81 | 29.34 | 27.54 | 31.71 | 28.66 | 23.67 | 19.89 | 22.03 |
| ScCM | 38.29 | 42.11 | | 21.26 | 22.81 | 22.67 | 22.09 | 25.71 | 24.73 | 25.68 |
| TaCM | 30.94 | 43.11 | 43.1 | | 64.07 | 45.83 | 45.24 | 24.43 | 19.13 | 21.74 |
| TvCM | 31.46 | 38.92 | 43.86 | 73.65 | | 47.27 | 45.45 | 23.7 | 20 | 20.99 |
| SbCM | 34.09 | 47.56 | 44.19 | 57.74 | 60 | | 60.25 | 22.94 | 19.77 | 20.22 |
| SpCM | 30.11 | 43.9 | 42.44 | 59.52 | 55.76 | 72.67 | | 24.12 | 20.9 | 22.47 |
| JbCM | 41.04 | 40.83 | 41.14 | 35.8 | 36.42 | 38.82 | 39.41 | | 68.75 | 67.7 |
| DsCM | 38.33 | 35.8 | 38.46 | 32.79 | 33.89 | 37.29 | 35.03 | 76.25 | | 65.22 |
| MpCM | 38.12 | 36.72 | 40.44 | 34.24 | 34.25 | 35.39 | 35.39 | 78.88 | 78.26 | |

**(B)**

| Identity \ Similarity | PaCDT | AfCDT | ScCDT | TaCDT | TvCDT | SbCDT | SpCDT | JbCDT | DsCDT | MpCDT |
|---|---|---|---|---|---|---|---|---|---|---|
| PaCDT | | 43.03 | 43.72 | 43.09 | 41.53 | 44.72 | 44.72 | 59.26 | 60.91 | 60.49 |
| AfCDT | 56.15 | | 38.87 | 39 | 39.09 | 39.42 | 38.59 | 44.12 | 44.12 | 46.64 |
| ScCDT | 59.11 | 56.68 | | 35.08 | 37.05 | 37.5 | 37.5 | 46.94 | 44.31 | 46.34 |
| TaCDT | 59.35 | 55.19 | 51.61 | | 74.58 | 61.3 | 64.35 | 43.7 | 42.86 | 40.76 |
| TvCDT | 60.08 | 56.38 | 53.39 | 86.44 | | 61.02 | 61.86 | 42.74 | 43.75 | 41.67 |
| SbCDT | 61.79 | 53.94 | 53.63 | 73.48 | 76.27 | | 71.74 | 40.76 | 40.34 | 41.6 |
| SpCDT | 59.35 | 55.6 | 54.44 | 74.78 | 75.42 | 81.3 | | 42.86 | 42.44 | 41.18 |
| JbCDT | 73.25 | 57.98 | 62.86 | 58.4 | 59.34 | 57.98 | 57.98 | | 76.69 | 73.31 |
| DsCDT | 73.25 | 58.82 | 61.79 | 58.4 | 59.17 | 56.3 | 56.72 | 87.29 | | 77.02 |
| MpCDT | 74.9 | 61.76 | 60.57 | 59.24 | 60 | 59.66 | 57.98 | 84.32 | 85.11 | |

**Figure S1. Sequence similarity and identity of the bifunctional enzymes.** Displayed are the percentages of sequence similarity (green) and identity (red) of the CM domain sequences (A) and CDT domain sequences (B) from all bifunctional fusion enzymes without their signal sequences. At the top left corners, the corresponding *MtCM and *PaCDT sequences are placed. Similarity and identity were calculated using the bioinformatic tools from P. Stothard, 2000.[9]

> *AfCMCDT-HC (GenBank accession no. **WP_083814300.1**)

CATATGCGTAAACCCCGCCACATTACAGCCTTGTTGTTTTGCCTGCTAACGTCACTTACAAGCGTGGCAGACA

ATCATTCGGAGCAGACGTTGTATCAATTGATGAGTGAAAGGCTCGCATTGATGCCCGAAGTGGCAAAATACAA

ATGGCATCACAACCTGCCAATCGAGGATTTAGCCCGCGAGGCGATGGTTCTGGAACGTACGGTATCTCGCACA

ACTGTATTAGATCCAATACATACGAAAACATTCTTCGGGCTGCAGATGACAGCTGCAAAGGCCATACAGGCAA

ATGTGTTTCAGTCACTAACTAACACAGATGTGGTTGCCTCCGACGTACGTTCTCTGAACGATGACCTGCGGCC

AAAATTGACCCTGCTCGGAGATCAGATAATAGAGCAGCTGCTTATTTCTTATCAAAATGGTACGCCTCTGAAC

AGAGCcCATTTCGATGCACATTTCGCGCACTTCGAACTTAATCCACAGATTAAAGACGGGCTCTTTAAGTCAC

TCGAACTTGTTCTTACCCCGCCCAACTTGGATCCTAGGGATACTCTCGCTAGATTAGAAAAAGATAAGACCCT

TCGGGTTGGTGTGACACTTGATTATGAACCGTTCTCTTATCAAGACAATGAGGGTAACAGAGCTGGTATAGAC

ATCGAGCTTGCGACCGCGCTGGCCAAAGAATTTGGGTATCGTATTGTGTGGGTTAAAACGTCATGGCCAACCC

TTATGGCAGATGCAGAAGATAACCTTTTTGACATTGCACTGTCAGGTATTAGTATCACCGCGCAACGTCAGCA

CCGCATGATGTTTAGCGCGCCATATCATACAGGAGGTAAAACAGCCATTGGACGCTGTTCATCAGTTGATGAG

TTAAACACACTAGCACTAATTGATCGGGCGGAAACCCGTATTATAGTTAACCCGGGAGGCACGAATGAACGGT

TTGTACGAAGTGCATTAACTAACGCATCTATACGTATCCACCCAGACAATAGAACAATATTTAACGAGCTTGT

TTCTGGGACTGCCGATGCCATGTTCACCGATAGCATAGAAGCCCAGTTGCAAGCGACAAAACATCCGTCATTG

TGCGTCCTGCTGGACCAACCTCTGACTTTTCAGCAAAAAGGTATACTGTTACAACCAGATCCGGAGTTAAAAA

AACGTATTGACACCTGGCTGCTCGATTATCTTAGCTCTCATGATGTTAGCGCTCTGTTCAGTAAACATGGCGT

TGACCCCGACCTCGAGCACCACCACCACCACCACTAATAATGACTAGT

> *ScCMCDT-HC (GenBank accession no. **WP_116808336.1**)

CATATGTCCCTGCACAAACGTATTTGTGTGGCACTTGCAATGCTCTTTACTGCTTCGGTGGCAGCGGCGGCGT

CATTTACAGGTCCCAACGAAGTCGCGCGCGTGTTTGATTTGATGCAACAGAGACTGGAACTGATGCGTGCTGT

TGCGGCTTGGAAGTATGCGAACAATGCCCCGGTTACCGACGCGGCGCGAGAACAGCAGGTGCTGGACGCCACT

GTTGCACAGGCCCAACGATTAGGAATTGATGCTGCTTCAGCCCGTGAACTGTTTGCACTCCAAATCCGGATGG

CCAGTGAGGTGCAGGAACATTTTATTGCGACTTGGCAGGCGCGTAAGTCTACAGACGAGGCAGTAAGAGATTT

ACAGCAAGAACTCCGTCCGCAGTTAGATCGTCTTGGTAACGAGCTGCTACACGCCATTTATTTAGCTCTGCCG

GAGCTGATGTCAGACGATTTTGCTGCCCGATATCAGTCGCAGGCAGCTAAAATAGCAATGCCTGGTCTGCGCC

AAGATGATCAGAGAGCTCTGCTGACAGCGGTTAGTAAGCTGCGTCCTGCCGCAATGCCAGCACGGGAACGCAT

TAAAGCGTCGAAAGTCTTACGTATTGGGATGACTGGTGATTACGCCCCGTTCACACTGGAGAGGGGTGGCGAG

CTTTCCGGCGCAGATGTTCAGATGGGAGAAGCCCTGGCGAAATCACTGGGTGCCCAACCACAATTTGTTTCTA

CTACTTGGTCCACCCTGATGCGTGATTATCAGGCAGGGCGCTTCGATGTTGCGCTGGGAGGTGTAAGCATAAC

CCCGGAGCGTACGAAAGTAGCAGCTTTTAGTGTGCCATATCATCAGGGCGGCAAAACGCCGATTGTGCGTTGT

GGTACAGAGAGTCGCTTCGACAGCGTGGAGGAGATTGATCGCCCAGATGTCCGCGTTGTAGTCAATCCCGGTG

GTACCAATCAGCAGTTCGTGCGAGAACGCCTGTCACACGCACATGTCACAGTCCATCCAGACAATCGGACAAT

CTTCGCAGAAATCGCCGGTGGACGCGCCGATGTTATGGTGACGGATGACGTAGAAGTAGATCTGCAGACGCGC

CGTGATAAGAGGCTTTGCCGGGCGACGTCAGCAACATTCACCCGGGGGGATAAAGCGATTTTGTTACCCCAGG

ACGAAGCGCTGAGGGGTCGTGTTGACCGCTGGTTACAGGGGCAAATTGCATCTGGAGCGGTCCAGGCTTGGTT

AGAATCAGCTTTGGCGGCGGAAGCACGTCTCCAGGCCGTCAATCTCGAGCACCACCACCACCACCACTAATAA

TGACTAGT

> *TaCMCDT-HC (GenBank accession no. **KJE41258.1**)

CATATGCTTCTGCTGATTAACCCCATCAAAATCATGAAGTTGCTGTCGAAAAAATCATATTTACTTTTGTTTA

TTTGTTTCTTTTCTCCATTTTCGGCGTTCGCGCAGACCCAAGCCAATGAACCCGGGAAATCATACCTCTACCA

ACTTATAAATTCACGTCTGGGCTACATGCAAGCCGTTGCTCTGTATAAATGGCAACATCAACGTGCGATTGAG

GACAGTGCTCGGGAACAGGTAGTAATTGAGAAAAGTGTTGCGAAAGCTATGGAACAAGGGCTGACGAGTGAAG

AGATTACACCCTTTTTTCAGATACAGATTACACTGGCGAAGAAAATTCAGGCATACTATCATAAACGCTGGTC

CGGCCATGGTGTGCCCACTCAGTTACTGTTGCCGGAAAACGCCCCCTCACTCGAAAAGATCCGCGCGGAGCTG

ATTTCTTTAGGTGCCGATATTATCACACGTCTAGCTGCAACCGATTCGTCTAAGGCATCACACGATTTCGAAC

AATTCAAGCAGGTTGTACAGCATGCTGCcCTAGACATTAACGATAAGGCGGCCCTGTTTAAAGCCCTGTCAAG

GATTAAACCACAACCGTACGCCAGCAGGCTGGACCGTATCTTATCGGAGAAAATTTTATATGTTGGTACAACC

GGCGATTATCGACCATTTAGCTTTTATGCAGATAACAAAAGAGCGGGCATTGATATAGTCCTTGCTCGCGATC

TGGCTCGGACGCTGGGTGCCTCGGCTGTATTTCTCCCGACTTCATGGCCGGGGCTGCTCGCCGATTTAGGTAC

GGGTCAATATGACATTATGATGAGTGGAATAAGTAAAAAACTGTTCCGTCAGCAGTTAGGGCTGTTTTCAGAT

AGCTACCATTCGGGTGGTAAAACCCCAATTAGCCTGTGTCGCAAAAAGCATCAATACAATTCGCTGGAAAAAA

TCGACCATCCGCAAACAAGACTGATCGTCAACAAAGGAGGTACAAACCAGCGCTTTGTCAATCAGCATATAAA

ACAGGCACAAGTTCTGGTACACGGAGACAACACAACCGTATTTGAGCAGATACTGGCGGGCCGTGCAGATGTT

ATGATCACTGATAAAATCGAAGTGGCAGTGCAAGCCAAAAATCACCCCCAGTTGTGTGGTACTATGAATGGTA

CGCTGAGTTATTCTGCTAAAGCcTTTCTTTTAGGACGGGACTTGATCTGGCTGGAATACGTAGACACATGGTT

GGAACAAGTTAAAAACGACGGTACGTTAAAACAGGTCTTTGAGCAGTATCTCGAGCACCACCACCACCACCAC

TAATAATGACTAGT

> SbCMCDT-HC (GenBank accession no. **ABN63218.1**)

CATATGATTTTGATGACATTTAGGTATTACTCGATGCGCTTGCACGCACTGTTGTGCCTAAGCGTGTTCGCCT

TACTATCGCTGCCAGCGCTAGCCGATACGGACGAAAACGCGGAGCTTTATGCAAATATGAACACTAGACTGAG

CTACATGCAACAGGTCGCACTTTACAAATGGCAGCATCAATTACCGATTGAAGACCTTGCGCGAGAAAAGATT

GTGCTGGCACAGAGTGTCACCGCTGCGGAATCACTCGGTATAACCAGCGTGGCGATCACAGATTTTTTTCAAG

TACAAATTGAACTGGCAAAGAAAATCCAGCGACAGTACCATCAACAGTGGCGCGAACATGGACTGCCTCAGAC

ACTCCAGACCAAACAAAACACTAAATTGAGTTTGGATAAAATCCGCCCAGCACTCACCACTCTGGGCCAAACG

ATTATTGAACAGATTGCAGAGCACCAGGATCAGCATGACTTCAGTGTATTTAATTTGGCGATTGACACTCCAC

TCGTATCGATAGAAGACAAGGCAGTCTTATTTCGCAGTCTTAGCTTGATAAAACCCAAAGTTTATCTTTCTAC

GCTGGATAAAATCATCGCTGAAAAAATTCTCTACGTCGGCACGACTGGCGACTACGAACCTTTCAGCTACTTT

GAGGCCGGTCAAATAAAGGGTATTGACATTGATCTGGCTAATAGACTGGCAGACTCGCTGGGTGCTCAGGCGG

TTTTTCTGCCCACTTCGTGGTCAAACCTCATTACCGATTTGTCATCAGAGCGATTTGACATCATGATGAGCGG

TATATCAAAGCAGCTGTTCCGCCAAAGAGTTGGGCTTCAGTCAGATATATATCTGGAAGATGGCAAAACCCCG

ATCAGCCTTTGTGCCAAAAAAGAGCGGTACGACAGCCTGGCAAAAATAGATAAACCTGATACCCGGATGATTG

TGAATAAGGGCGGCACGAACCAACGTTTCGTGGATGCTAACATCAAACAAGCAAAAATCACCGTTCATAGTAG

TAACGTGACCATTTTTCAGGAACTAATCGCGAACCGTGCTGATGTCATGATTACAGATCGGGTTGAAGTTCAG

TTGCAGACTAAAAAACATACTGAATTGTGTAGCACCATGCCGAACAATCTGAACTATAGTGCTAAAGCATTCC

TCATGGGTAGAGATCCGATTTGGAAAGAGTATGTCGACGCGTGGTTGGAACTGTCTATTAAAGATGGTAGCGT

GAGCAATATATTCAACCACTATATTCTCGAGCACCACCACCACCACCACTAATAATGACTAGT

> *JbCDTCM-HC (GenBank accession no. **ELX09769.1**)

CATATGCAACGGTTCATTCGACATAGCATGAGACAGATCGCGGTGCTGGGACTCTTAGCCGGTATGATGGCCT

CTGTTCAGGCCGGAGCGGATCATAGGCTGGATGACATCACGGCGCGCGGTGTCTTGCGGGTGGGTACTACTGG

CGATTATAAACCGTTTAGTTCTCGGGCAGGTAATGACTTTGTTGGACTGGATATCGAGCTTGCAGCGGACCTG

GCCCGTACGCTGGGCGTCCCGGTGCAGATTGTGCCGACTTCCTGGCCTACACTTATGAAAGACTTTGGCGATG

GGAAATTCGACATCGCACTGGGCGGTGTTAGCATTACCCCCGAGCGGCAGAAGCAGGGTTTGTTTTCAGTTAG

CTATCTGCGGGATGGAAAAACACCCATTACTCGATGTGAGAACTCAGCACGGTTTCAAACGTTGGCACAGATC

GACCAACCGGGCGTAAGGCTGGTGGTGAATCCAGGCGGGACTAACGAGCGTTTTGCTCGCTCGCAGGCGCCAA

ATGCCCAACTCACCGTCTACCCTGATAATGTGACCATTTTCGATCAGATTGTAACGGGCGCAGCAGATTTGAT

GATCACGGATGCAATTGAAACGCGGCTGCAACAACGACTTCGCCCTCAGCTTTGCGCAGTACATCCAGATACA

CCTTTTGACTTCGCCGAAAAAGCGATTCTCTTGCCCCGGGATGTTGCGTTCAAGGCAGTAGTCGATAAATGGC

TTCAACAGAGGATTGCATCAGGGGCTGTACAGCGGAGCGTTGATCGGTGGCTGGATTTTCCCTGGGGCTTGGA

ACCCTTACGTTTGGCCATTGACCAGCGGCTGCTGTTGGCTCAGGCCGTTGCCCGTGCTAAATGGAACGTGCAG

GCTCCGATTGAAGATCTTGGGCGGGAAGCCCAAGTGATACAGGCGGCTGTCAAAGAAGGCGCTGCACTGGGTC

TGCCGAAGGTTTGGATTGAAACTGTATTTCGTGCACAGATTGAAGCAAGCAAAACCGTGCAACGCGAACTGTT

CGCCCAGTGGTCAGCCCAACAGGCGGGCAAATTTGATGACGCCCCTGACTTAGCAAAGACCATCCGTCCGGAA

CTAGACCGCTTAACTACTCAGCTGTTACGTTCGATGGCATCGAATCAGACTGTGTTAAACGATGAGGCTCGTA

AAGCAGATGTAGCGCGTGCAATGCGGGCTTTAGAAGCCAGAGCTTTATCTCCTCAAGCGGCGACCCAGGCTCT

CGCACCGTTTTTCCTCGAGCACCACCACCACCACCACTAATAATGACTAGT

> *DsCDTCM-HC (GenBank accession no. **WP_072786685.1**)

CATATGAAGAGATTGCTATTGAGCACTCTGCTGGTTACGGCTTTAGCGAGTGCACATGCAGGCCGTTTGGAAG

AGATTCATGCCCGTGGAGTTTTAAGGGTGGGTAGCACCGGGGACTATAAACCATTCAGCTATCGTGCAGGTGC

GAATGATTTTATTGGGCTGGATGTCGAGCAGGCCGGTGAATTAGCTCGCGCTATGGGTGTTAAACTGGAAATC

GTGCCGACAAGCTGGCCCACGCTGATGACGGACTTTGGCGCGGACAAATTTGATATTGTACTGAGTGGTGTGT

CGGTGACTGCAGAACGTCAACAACAGGCTCTATTTTCAGTCAGTTACCTACGTGATGGCAAAACGCCAATTAC

TCGGTGTGAGAATCAACTGCGTTTTCAGACGTTGGAGCAGATCAATCAGCCTGCAGTACGTCTCATTGTCAAT

CCTGGAGGTACTAATGAACGATTCGCTCGTGCTCATGCACCGCATGCTCAGTTGACGGTATACCCTGACAACG

TTACAATTTTTGGCCAGATTGTTTCCGGTGCCGCGGATTTAATGATGACTGATGCCATTGAAACTCGCCTGCA

GCAGCGTTTGCATCCAGAATTGTGTGCTGTTCACCCCGATGCCCCGTTTGATACAGCCGAAAAGGCAATATTA

CTGCCGCGTGATGCAGAACTGAAAATATATGTGGATACTTGGCTCCAACAGCGAATTAGCTCTGGTGGCCTCC

AGAAATCCTTTGATCGGTGGTTGGATTATCCATGGGCGCTTGAGCCTTTACGCCAGGCCATCGATGAAAGACT

GCTTCTGGCCGAAGCGGTGGCCAGGGCTAAGTGGAATGTGCAAGCTCCAATTGAAGATCTGCCTCGTGAGGCT

CAGGTAATAGCCGCGGCCGTACAGCAAGGCCGTACACTGGGATTACCCGACGCTTGGGTTGCAGCCGTTTTTA

AGGCCCAGATAGAAGCTAGCAAAACTGTGCAACGCGAGTTGTACGCGAAATGGAAGGCACAGCAGGCAGGGCA

CTTTGATGATGCGCCGGACCTGGCAAATACGATACGCCCGCAACTTGACCGTATCACGACCCAGTTACTGAGA

GCAATGGCTGATAATCAGGCGACATTAAAAGATACTGCAAGATTAATCCGTCCTCTGGAGGCCGCCGCCCTGT

CCCCTGCGGCGGCGGCGCAAGCCCTTGCTCCATTAAGCGCGCACGTAGTCGTCCGCTTTCTCGAGCACCACCA

CCACCACCACTAATAATGACTAGT

> *MpCDTCM-HC (GenBank accession no. **PQP01982.1**)

CATATGTTCGGCCGCGTCTGGATGCGTCCGGTGGCATCGGCGGTCATGCTGGCAGCCGCGCTGGCGCCGGCTC

AGGCAGGTCATTTAGATGATATTGCCGCGCGTGGAGTGCTCCGCGTAGGCTCAACAGGCGATTACAAGCCTTT

```
TTCCTACCGTCAGACCGATGGTGGTTTTATTGGTATGGATGTTGACCTTGCAGGTGAGCTGGCCCGCTCTCTC
GGCGTTCGCCTGGAGCTTGTTCCAACCACATGGCCCACGTTAATGGCGGATTTGGGTGCCGGTAAATTCGATC
TTGCACTATCCGGTGTCAGCGTCACAGCAGAGCGTCAACGCCAGGCCCTGTTTAGTGTGCCGTATCTCCATGA
TGGCAAAACACCAATTACAAGGTGCGAAAATGTGGCCCGTTTCCAGACCCTTGCACAGATCGATAGGCCTGAA
GTGCGTCTGATCGTGAACCCTGGAGGGACAAATGAGCGGTTCGCGCGAGCTCAAGCCCCCCGGGCGCGGCTGA
TTGTATACCCGGACAATGTTACCATATTCGGTCAGATCGTATCAGGAGCCGCCGACCTGATGATGACCGATGC
AATAGAAACACGTTTGCAGCAGAGATTGCATCCACAACTATGCGCTGTGCACCCAGAAGCCCCCTTTGATATG
GCAGATAAAGCGATCTTACTGCCGCGGGATCCGGCGCTCAAGACAGTGGTGGACCGATGGTTGCAGCAGCGGT
TAGATAATGGGGATGTGCCAAAGCGGTTAGATCGCTGGCTCGCCTTCCCGTGGGGGTTGGAACCCTTACGTCA
AGCCATAGATCAGCGATTACTACTTGCACAAGAAGTTGCTCGCGCAAAATGGAACGCTAAAGCGGCTATAGAA
GATCTACCACGCGAAGAACAGGTGATTGCAGCCGCAGTTCGACAGGGCAGTGCTCTAGGTTTGCCAGAAGCAT
GGGTGCGCACAGTCTTCCGTGCTCAGATTGAAGCAAGCAAAACAGTACAGCGCGCCCTATACGGCCGTTGGCA
GGCTGAGGGCGCTGGGAGATTTGATGATGCTCCGGATTTGGCAGGGTCAGTCAGGCCAGAACTGGACCGTCTT
ACTACACAACTGCTGCGGGCCATGGCAGATAATCAAGCTCTGTTACATGATGCCGACCGAAAAGCTGATATAG
CTGTAGTCATGCATGCTCTGCAGGCTCATGCCGTGAATCCCGCGGCCGCGGGTCAAGCATTGGCGCCCTTTCT
GGCTAGTGCACCGAGCGCTGGGGAACTCGAGCACCACCACCACCACCACTAATAATGACTAGT
```

**Figure S2. Codon-optimized nucleotide sequences for expression in *E. coli* of the genes for the bifunctional fusion enzymes with an appended C-terminal His-tag.** The sequences were derived by codon-adapted reverse translation from the listed GenBank accession numbers. Note that **KJE41258.1** (encoding *TaCMCDT) was subsequently updated to **WP_160298287.1**, which specifies a protein with an 11 residue shorter signal sequence. This update does not affect this work, however, since all experiments were carried out with the mature protein, of which the sequence beyond the signal peptide is identical for both accession numbers.

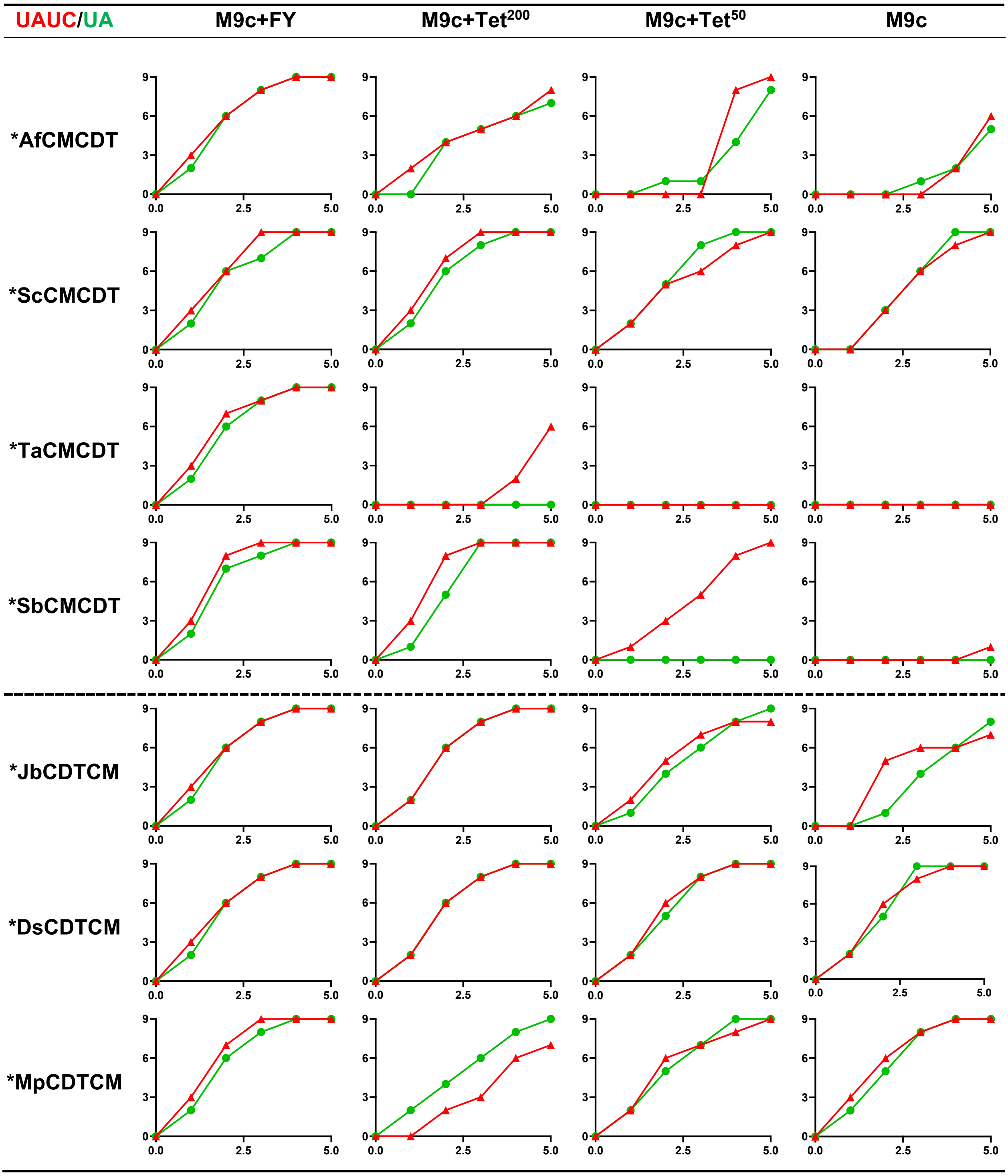


**Figure S3.** ***In vivo* complementation assays.** Shown are the growth curves of *E. coli* strains KA12/pKIMP-UAUC (UAUC, red triangles) and KA12/pKIMP-UA (UA, green dots) transformed with expression plasmid pKTCTET that encodes the specified cytoplasmic versions of the bifunctional fusion enzyme, indicating its efficiency in CM complementation or CM+CDT complementation, respectively. The clones were grown on M9c minimal agar plates with supplementation of Phe and Tyr (FY, viability control), addition of 200 or 50 ng/mL tetracycline (Tet$^{200}$ or Tet$^{50}$), or no additions, while scoring the size of growing colonies from 0 (no trace of growth) to 9 (ø > 3 mm) (y-axis) over 5 days at 30°C (x-axis).

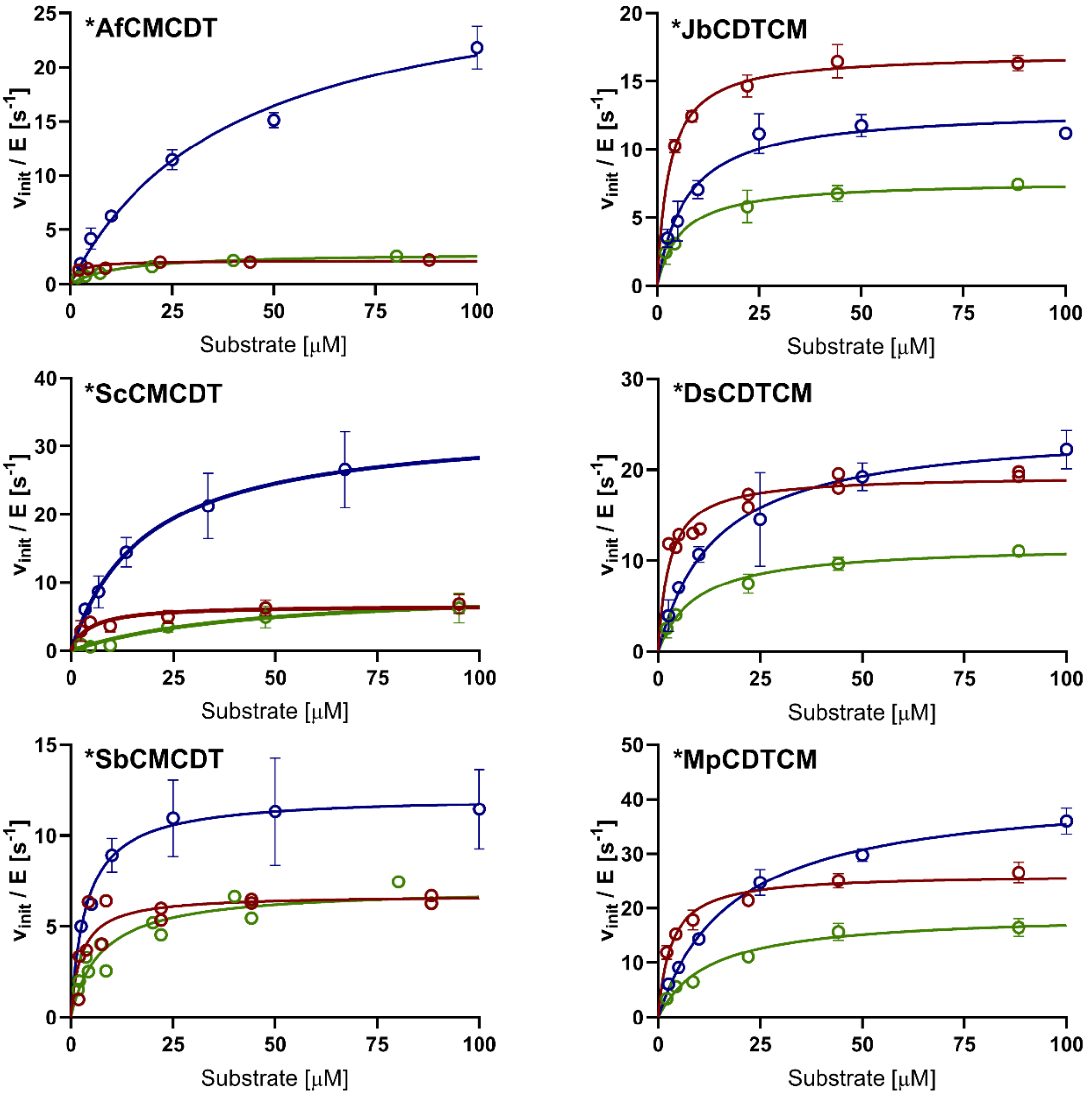


**Figure S4. Michaelis-Menten plots of the bifunctional fusion enzymes.** Shown are the Michaelis-Menten plots of CM (red), CDT (blue), and coupled CM+CDT assays (green) of six bifunctional enzymes. Two independently prepared biological replicates were measured in each case. The curves were fitted to the mean activity values (each data point shown with standard deviation bars; assuming identical experimental substrate concentrations).

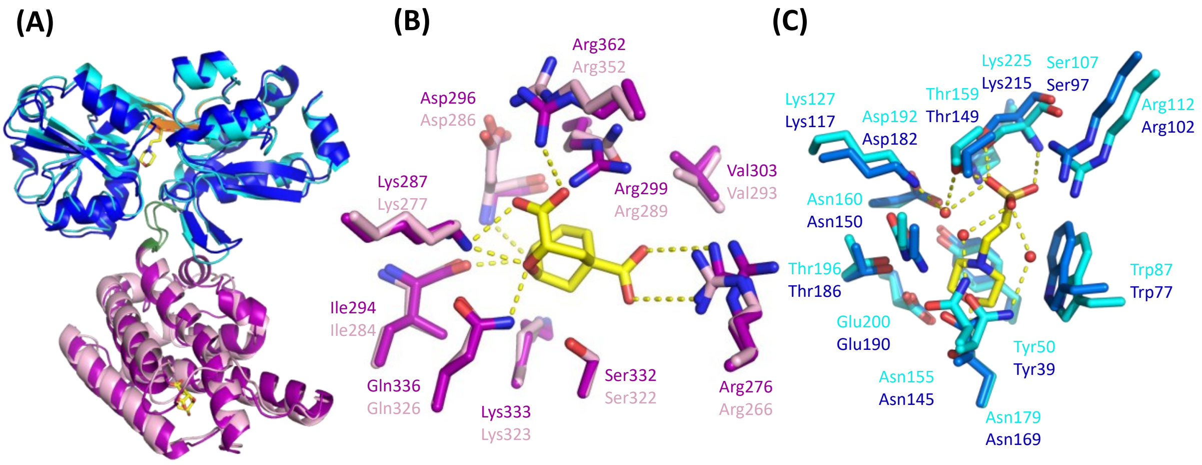


**Figure S5. Structural alignment of *JbCDTCM and *DsCDTCM.** (A) Superimposition of *JbCDTCM (PDB ID: 8CQ4, this work; cyan for *CDT, and magenta for *CM domain) and *DsCDTCM (PDB ID: 8CQ6, this work; dark blue for *CDT, and pink for *CM domain). The two enzymes adopt very similar structures, with r.m.s.d. = 1.4 Å ($C_\alpha$ atoms). (B) Superimposition of active sites of the *CM domains of *JbCDTCM and *DsCDTCM (r.m.s.d. = 0.4 Å; all atoms). A TSA molecule (yellow carbons) is superimposed from the *MtCM structure (PDB ID: 2FP2)[10]. Hydrogen bonds are shown with dashed yellow lines. (C) Superimposition of active sites of the *CDT domains of *JbCDTCM (complex with MES, yellow carbons) and *DsCDTCM, with r.m.s.d = 0.9 Å (all atoms).

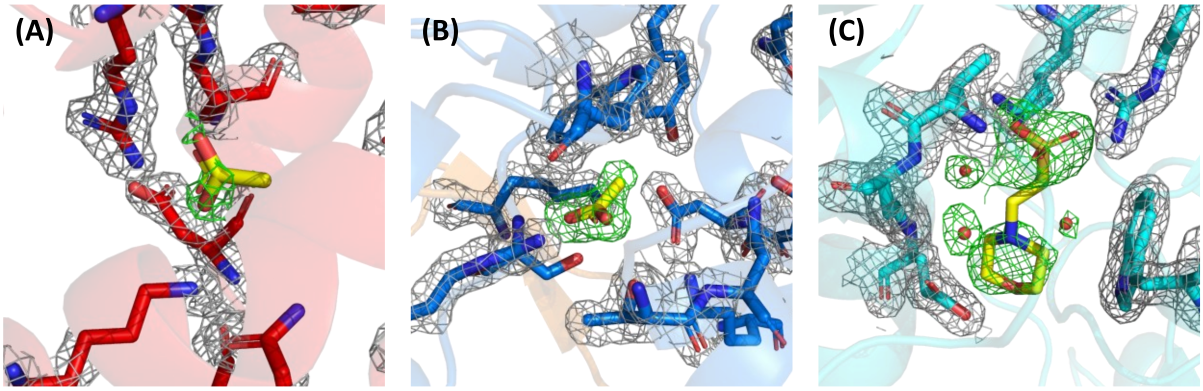


**Figure S6. Ligands in active sites of the bifunctional enzymes.** (A) Acetate bound to the CM active site of *AfCMCDT (PDB ID: 8CQ3, this work), shown as yellow sticks, with CM active site residues rendered as red sticks. (B) Acetate bound to the CDT active site of *AfCMCDT, shown as yellow sticks. CDT active site residues are depicted as blue sticks. (C) 2-(*N*-morpholino)ethanesulfonic acid (MES, yellow sticks) bound to the *CDT active site of *JbCDTCM (PDB ID: 8CQ4, this work). Active site residues are shown as cyan sticks. The $\sigma_A$-weighted m*F*o-D*F*c difference electron density maps for the ligands are shown at 3.0 σ (green mesh); $\sigma_A$-weighted 2m*F*o-D*F*c maps are depicted at 1.5 σ (grey mesh).

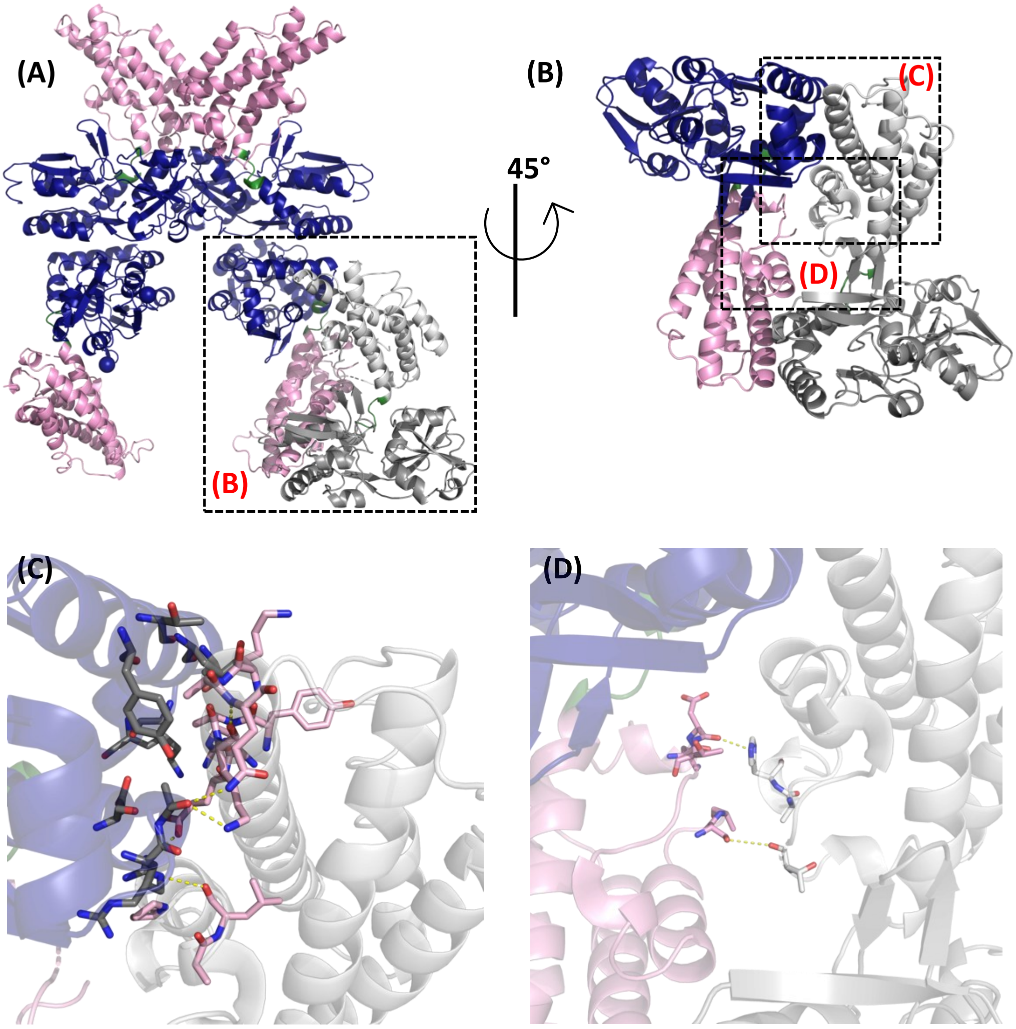


**Figure S7. Crystal structure of *DsCDTCM.** (A) Asymmetric unit (a.u.) content of *DsCDTCM crystal (PDB ID: 8CQ6, this work), colored in pink (CM) and dark blue (CDT). The chain in dark/light grey, showcasing the *DsCDTM dimer, is reconstructed by crystallographic symmetry. (B) Head-to-tail dimer of *DsCDTCM reconstructed by crystallographic symmetry and outlined by the box in (A). Two different dimerization interfaces are highlighted in boxes. (C) Dimerization interface between CM and CDT domains from two different chains. (D) Dimerization interface between two CM domains from two different chains.

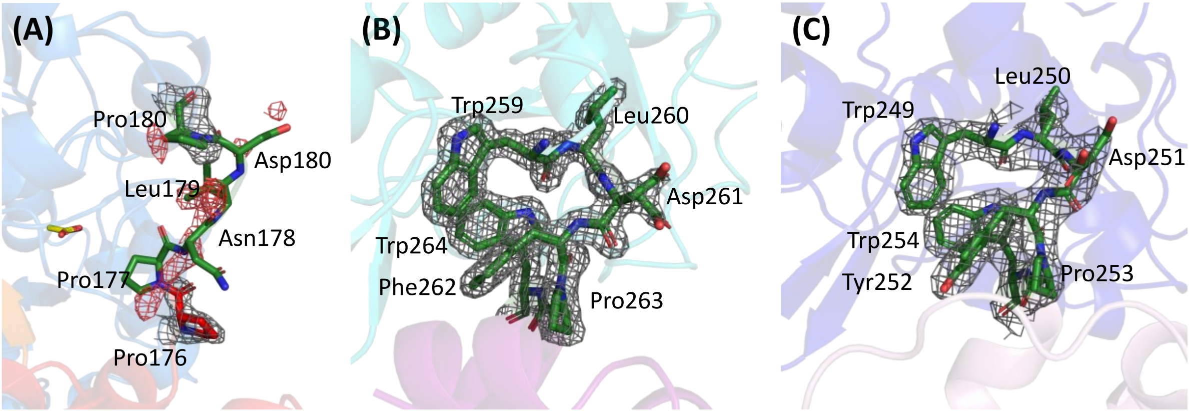


**Figure S8. Electron density maps of the linker connecting the enzyme domains of *AfCMCDT, *JbCDTCM, and *DsCDTCM.** Shown are the protein crystal structures determined in this work, *i.e.* (**A**) *AfCMCDT (PDB ID: 8CQ3), (**B**) *JbCDTCM (PDB ID: 8CQ4), and (**C**) *DsCDTCM (PDB ID: 8CQ6), with a focus on the linker region (green sticks between the CM and CDT domains). $\sigma_A$-weighted 2m***Fo***-D***Fc*** maps are shown as grey mesh at 1.5 σ. The $\sigma_A$-weighted m*Fo*-D*Fc* difference density map for the poorly resolved residues (P156-P160) in *AfCMCDT is depicted at 3.0 σ as red mesh.

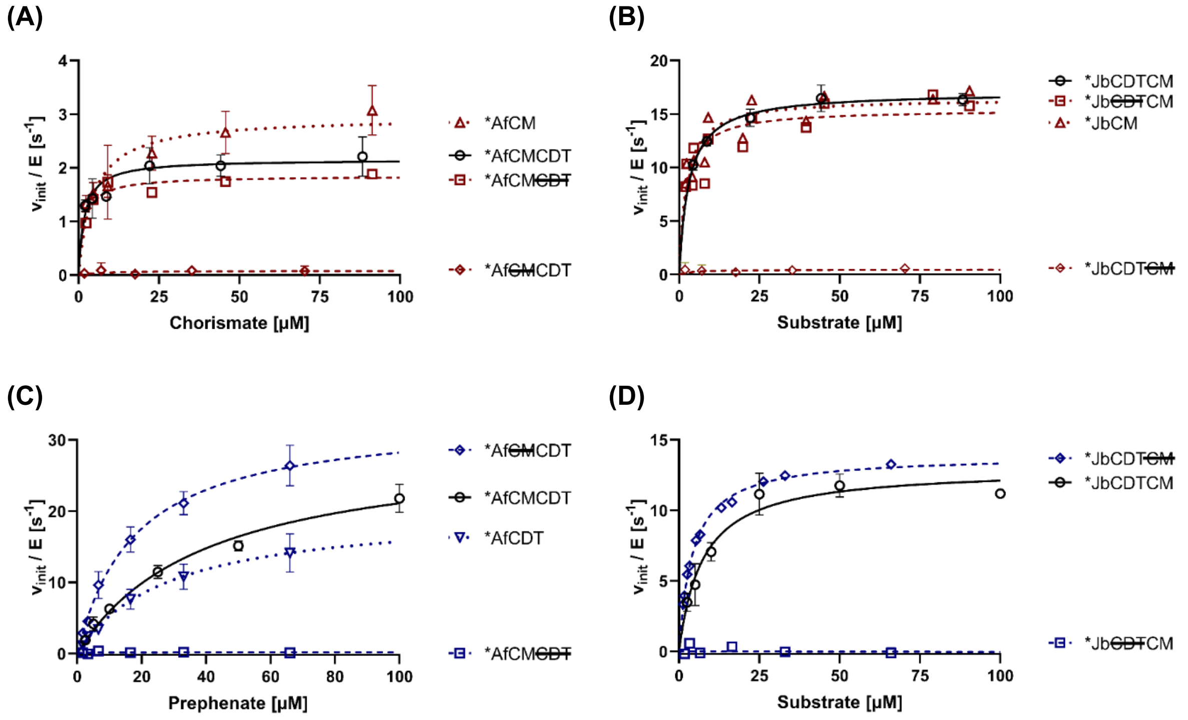


**Figure S9. Michaelis-Menten plots of active site KO and single split domain variants of *AfCMCDT and *JbCDTCM.** CM activity (A) and (B), as well as CDT activity (C) and (D) of the different formats of *AfCMCDT and*JbCDTCM, respectively, is displayed. Two independently prepared biological replicates were measured of each variant and the curve fitted to the calculated mean at each substrate concentration. Standard deviation bars are given for identical substrate concentrations in the measurement of the two replicates. The wild-type data are taken from Figure S4.

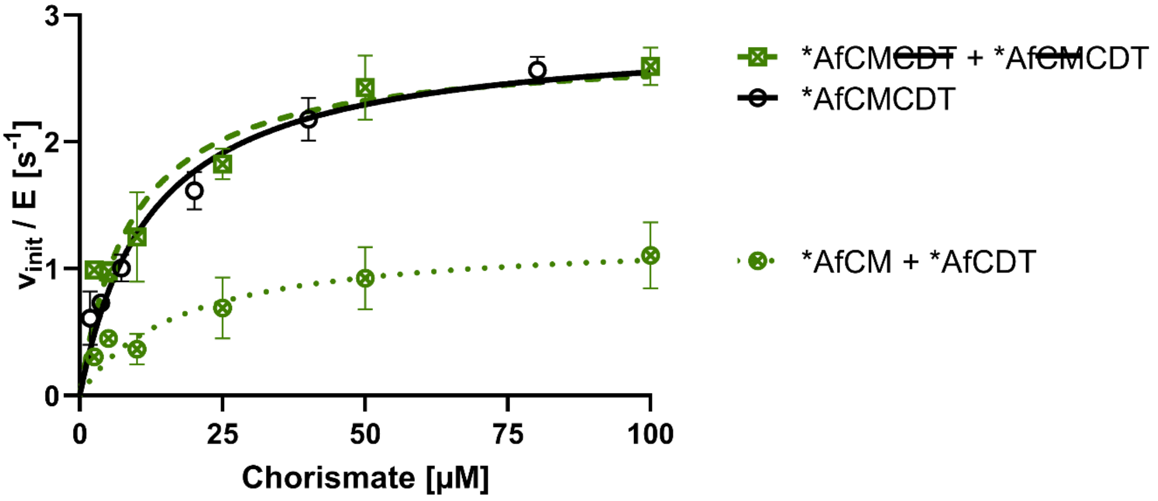


**Figure S10. Catalysis of the sequential CM + CDT reaction of mixed active site KO or split domain variants of *AfCMCDT compared to the parental bifunctional fusion enzyme.** Shown are Michaelis-Menten plots of coupled CM+CDT kinetic measurements with equimolar concentrations of either *AfCM~~CDT~~ and *Af~~CM~~CDT active site KO variants (green dashed line) or *AfCM and *AfCDT single split domains (green dotted line). The wild-type *AfCMCDT curve is taken from Figure S4 (black solid line). Equimolar concentration of CM and CDT active sites in each assay allow for direct comparability with the wild-type *AfCMCDT activity. For each data point, two independently prepared biological replicates were averaged with the bars indicating the standard deviation. The curves were fitted to the calculated mean at each substrate concentration using the Michaelis-Menten equation.

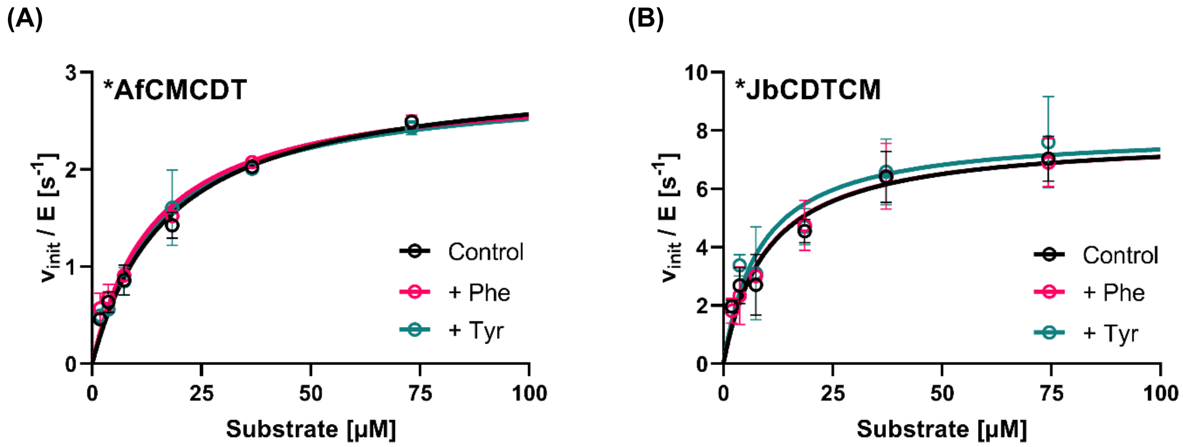


**Figure S11. Testing for feedback regulation of CM or CDT activity by Phe or Tyr.** Michaelis-Menten plots of coupled CM+CDT kinetic assays with (A) *AfCMCDT and (B) *JbCDTCM in the presence of 200 μM L-Phe (red curve) or 200 μM L-Tyr (green curve), corresponding to an 80,000-fold molar excess over the enzyme concentration, in comparison to a control assay in the absence of L-Phe or L-Tyr (black curve, data from the corresponding plot of Figure S4). The curves were fitted to the mean activity values of two independent biological replicates at a particular substrate concentration, with error bars depicting standard deviations.

***AfCMCDT**
**WP_083814300.1**

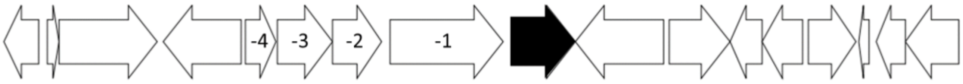


| | | |
|---|---|---|
| -4 | TetR_N | Bacterial regulatory proteins, *tetR* family |
| -3 | Glyoxalase | Glyoxalase/bleomycin resistance protein/dioxygenase superfamily |
| -2 | FAA_hydrolase | Fumarylacetoacetate (FAA) hydrolase family |
| -1 | Peroxidase | Peroxidase |

***ScCMCDT**
**WP_116808336.1**

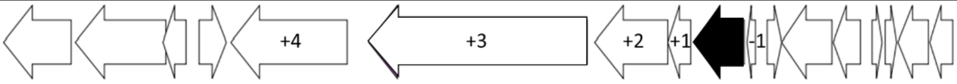


| | | |
|---|---|---|
| -1 | unknown | - |
| +1 | LemA | LemA family |
| +2 | Peptidase_M48 | Peptidase family M48 |
| +3 | ABC_tran | ABC transporter |
| +4 | TonB_dep_Rec | TonB dependent receptor |

***TaCMCDT**
**WP_160298287.1**

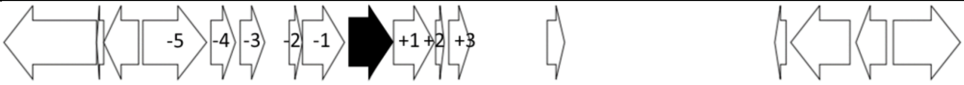


| | | |
|---|---|---|
| -5 | Trans_reg_C | Transcriptional regulatory protein, C terminal |
| -4 | SBP_bac_3 | Bacterial extracellular solute-binding proteins, family 3 |
| -3 | AAA_5 | AAA domain (dynein-related subfamily) |
| -2 | Cytochrom_C | Cytochrome c |
| -1 | COX1 | Cytochrome c and quinol oxidase polypeptide I |
| +1 | ATPase | KaiC |
| +2 | DUF1330 | Domain of unknown function (DUF1330) |
| +3 | SBP_bac_3 | Bacterial extracellular solute-binding proteins, family 3 |

***TvCMCDT**
**WP_053046572.1**

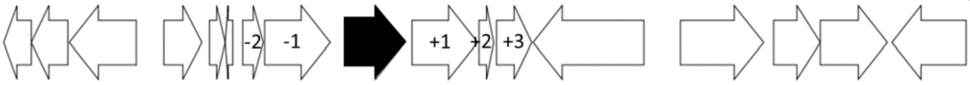


| | | |
|---|---|---|
| -2 | Cytochrom_C | Cytochrome c |
| -1 | COX1 | Cytochrome c and quinol oxidase polypeptide I |
| +1 | ATPase | KaiC |
| +2 | DUF1330 | Domain of unknown function (DUF1330) |
| +3 | SBP_bac_3 | Bacterial extracellular solute-binding proteins, family 3 |

***SbCMCDT**
**ABN63218.1**

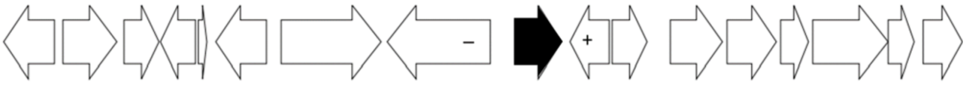


| | | |
|---|---|---|
| – | Peptidase_M16_C | Peptidase M16 inactive domain |
| + | Asparaginase | Asparaginase, N-terminal |

***SpCMCDT**
**WP_077752318.1**

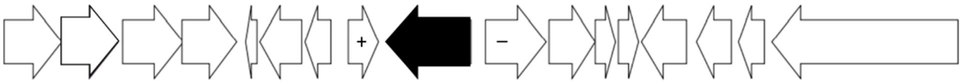


| | | |
|---|---|---|
| – | AraC_binding | AraC-like ligand binding domain |
| + | unknown | - |

***JbCDTCM**
**ELX09769.1**

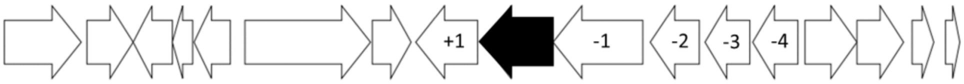


| | | |
|---|---|---|
| -4 | SIMPL | Protein of unknown function (DUF541) |
| -3 | SIMPL | Protein of unknown function (DUF541) |
| -2 | CDP-OH_P_transf | CDP-alcohol phosphatidyltransferase |
| -1 | HMGL-like | HMGL-like (pyruvate carboxylase) |
| +1 | Lumazine_bd_2 | Putative lumazine-binding |

***DsCDTCM**
**WP_072786685.1**

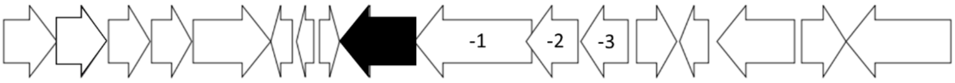


| | | |
|---|---|---|
| -3 | MipA | MltA-interacting protein MipA |
| -2 | CPBP | CPBP intramembrane metalloprotease |
| -1 | AI-2E_transport | AI-2E family transporter |

***MpCDTCM**
**PQP01982.1**

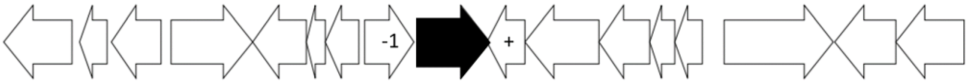


| | | |
|---|---|---|
| -1 | LysR_substrate | LysR substrate binding domain |
| + | GyrI-like | GyrI-like small molecule binding domain |

**Figure S12 Genomic neighborhood analysis of the exported bifunctional fusion enzyme genes.** Listed is a selection of the RODEO[11] output using the exported bifunctional fusion enzymes as query input (protein accession numbers in bold and underlined). The arrows in the graphs indicate the orientation and relative size of the genes surrounding the open reading frame of the exported bifunctional fusion enzyme (black arrow). The numbers in the arrows serve as reference to the list with the corresponding protein families and a short description. Genes that have the same orientation as the exported bifunctional fusion enzymes are potentially in the same operon. The exported bifunctional fusion enzymes themselves belong to the 'SBP_bac_3' protein family (Bacterial extracellular solute-binding proteins, family 3).

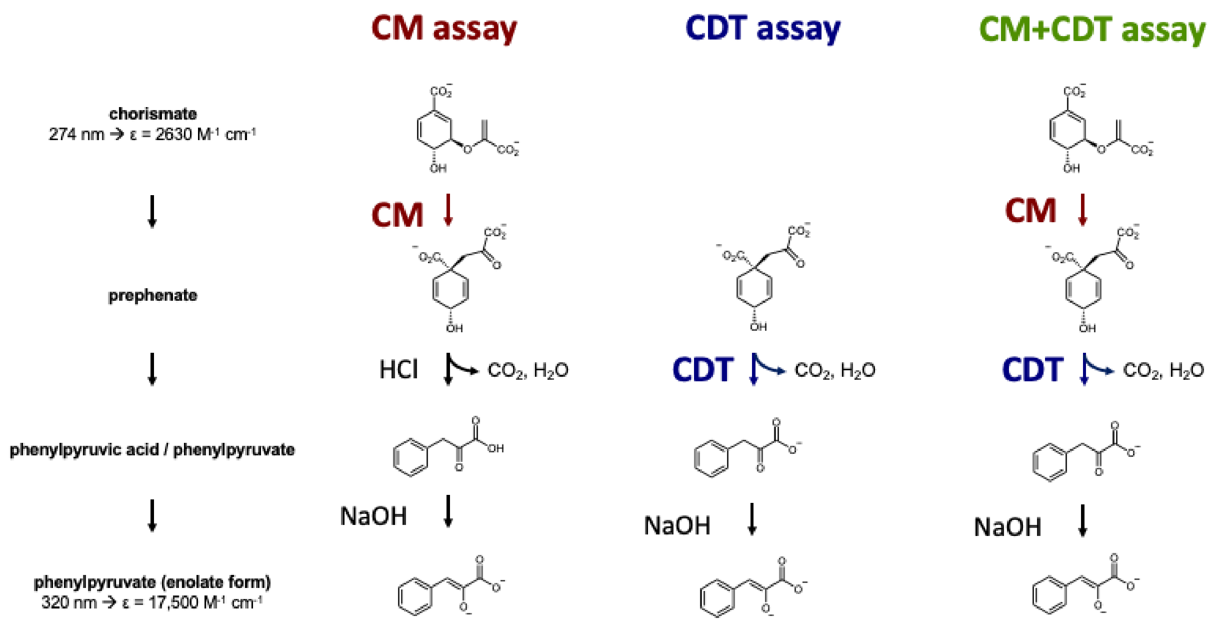


**Figure S13. Illustration of reactions during the *in vitro* discontinuous kinetic assays.** The reaction pathways for the CM (red), CDT (blue) and coupled CM+CDT (green) assay are shown. Chorismate is the substrate for CM and CM+CDT assays, where it is enzymatically converted to prephenate. Prephenate in turn gets converted by chemical decarboxylation and dehydration upon acidification with HCl or by CDT to phenylpyruvic acid or phenylpyruvate, respectively. In the CDT assay, prephenate is the added substrate and is incubated for enzymatic conversion to phenylpyruvate, thus no acidification step is required. In all three assays, NaOH is added in the final step to shift the pH to the alkaline range resulting in the formation of the enolate form of phenylpyruvate, which exhibits a high extinction coefficient (ε) of 17,500 $M^{-1}$ $cm^{-1}$ at 320 nm.

### Supplementary Tables

#### Table S1 Data processing and anisotropy statistics from *STARANISO*[12]

| | *AfCMCDT | *JbCDTCM, MES | *DsCDTCM |
|---|---|---|---|
| Resolution range | 52.5-1.7 (1.73-1.55) | 65.6-1.65 (1.83-1.65)[a] | 99.9-2.44 (2.66-2.44) |
| Diffraction limit #1 (Å) | 2.08 | 1.95 | 3.71 |
| Principal axes (orthogonal basis) | 0.9777, 0.0000, -0.21000 | 1.0000, 0.0000, 1.0000 | 1.0000, 0.0000, 0.0000 |
| Principal axes (reciprocal lattice) | 0.784 * - 0.621 c* | a* | a* |
| Diffraction limit #2 (Å) | 1.67 | 1.81 | 2.54 |
| Principal axes (orthogonal basis) | 0.0000, 1.0000, 0.0000 | 0.0000, 1.0000, 0.0000 | 0.0000, 1.0000, 0.0000 |
| Principal axes (reciprocal lattice) | b* | b* | b* |
| Diffraction limit #3 (Å) | 1.55 | 1.65 | 2.44 |
| Principal axes (orthogonal basis) | 0.2100, 0.0000, 0.97777 | 0.0000, 0.0000, 1.0000 | 0.0000, 0.0000, 1.0000 |
| Principal axes (reciprocal lattice) | 0.159 a* + 0.987 c* | c* | c* |
| Wilson *B*-factors (Å$^2$) | | | |
| Eigenvalue #1 (Å) | 46.6 | 35.3 | 135.41 |
| Principal axes (orthogonal basis) | 0.9900, 0.0000, -0.1408 | 1.0000, 0.0000, 0.0000 | 1.0000, 0.0000, 0.0000 |
| Principal axes (reciprocal lattice) | 0.824 a* - 0.567 c* | a* | a* |
| Eigenvalue #2 (Å) | 18.3 | 31.2 | 52.8 |
| Principal axes (orthogonal basis) | 0.0000, 1.0000, 0.0000 | 0.0000, 1.0000, 0.0000 | 0.0000, 1.0000, 0.0000 |
| Principal axes (reciprocal lattice) | b* | b* | b* |
| Eigenvalue #3 (Å) | 17.2 | 23.6 | 47.1 |
| Principal axes (orthogonal basis) | 0.1408, 0.0000, 0.9900 | 0.0000, 0.0000, 1.000 | 0.0000, 0.0000, 1.0000 |
| Principal axes (reciprocal lattice) | 0.103 a* + 0.995 c* | c* | c* |

Diffraction limits and eigenvalues of overall anisotropy tensor on |F|s are displayed alongside the corresponding principal axes of the ellipsoid fitted to the diffraction cut-off surface as direction cosines in the orthogonal basis and in terms of reciprocal unit-cell vectors.

**Table S2 Calculated and observed molecular masses of the produced enzyme variants**

| Wild-type Enzyme | $M_{\mathrm{r(calc)}}$ [Da][a] | $M_{\mathrm{r(obs)}}$ [Da] | Enzyme variants | $M_{\mathrm{r(calc)}}$ [Da][a] | $M_{\mathrm{r(obs)}}$ [Da] |
|---|---|---|---|---|---|
| *AfCMCDT | 45467.1 | 45466.8 | *Af~~CM~~CDT | 45410.0 | 45409.3 |
| *ScCMCDT | 45825.5 | 45694.7[b] | *AfCM~~CDT~~ | 45466.2 | 45466.4 |
| *TaCMCDT | 45623.8 | 45624.3 | *AfCM | 19045.5 | 19044.8 |
| *SbCMCDT | 45535.6 | 45536.8 | *AfCDT | 27624.9 | 27624.2 |
| *JbCDTCM | 43994.8 | 43995.5 | *JbCDT~~CM~~ | 43939.7 | 43940.5 |
| *DsCDTCM | 44157.9 | 44026.0[b] | *Jb~~CDT~~CM | 43993.8 | 43994.2 |
| *MpCDTCM | 44530.4 | 44399.7[b] | *JbCM | 18008.5 | 17876.7[b]/18008.7 |
| | | | *JbCDT[c] | - | - |

[a] The $M_{\mathrm{r(calc)}}$ was corrected for the expected disulfide bond formation in the CDT domains (-2 Da) and a directly attached N or C-terminal $His_6$-tag (+822.9 Da).

[b] The difference between observed and calculated molecular mass agrees with cleavage of the N-terminal Met residue (-131.0 Da).

[c] No soluble protein was obtained for variant *JbCDT.

### Table S3  Reaction composition and timing for the three discontinuous kinetic assays

| # | Substrate | Enzyme | reaction time at 30°C | HCl (2 M) | NaOH (10/5 M) |
|---|---|---|---|---|---|
| | µM | nM | min | µL | µL |
| A1 | 2.5 | […] | 0 | 100/0 | 100/200 |
| A2 | | | 0.25 | 100/0 | 100/200 |
| A3 | | | 0.5 | 100/0 | 100/200 |
| A4 | | | 0.75 | 100/0 | 100/200 |
| B1 | 5 | […] | 0 | 100/0 | 100/200 |
| B2 | | | 0.5 | 100/0 | 100/200 |
| B3 | | | 0.75 | 100/0 | 100/200 |
| B4 | | | 1 | 100/0 | 100/200 |
| C1 | 10 | […] | 0 | 100/0 | 100/200 |
| C2 | | | 0.5 | 100/0 | 100/200 |
| C3 | | | 0.75 | 100/0 | 100/200 |
| C4 | | | 1 | 100/0 | 100/200 |
| D1 | 25 | […] | 0 | 100/0 | 100/200 |
| D2 | | | 0.5 | 100/0 | 100/200 |
| D3 | | | 1 | 100/0 | 100/200 |
| D4 | | | 2 | 100/0 | 100/200 |
| E1 | 50 | […] | 0 | 100/0 | 100/200 |
| E2 | | | 1 | 100/0 | 100/200 |
| E3 | | | 2 | 100/0 | 100/200 |
| E4 | | | 4 | 100/0 | 100/200 |
| F1 | 100 | […] | 0 | 100/0 | 100/200 |
| F2 | | | 1 | 100/0 | 100/200 |
| F3 | | | 2 | 100/0 | 100/200 |
| F4 | | | 4 | 100/0 | 100/200 |

| […] | CM | CDT | CM+CDT |
|---|---|---|---|
| Variant | nM | nM | nM |
| *AfCMCDT | 10 | 5.0 | 10.0 |
| *AfCM~~CDT~~ | 10 | 2.5 | – |
| *Af~~CM~~CDT | 10 | 2.5 | – |
| *AfCM | 10 | n.p. | – |
| *AfCDT | n.p. | 2.5 | – |
| *ScCMCDT | 2.5 | 2.5 | 2.5 |
| *TaCMCDT | 10.0 | 100.0 | 100.0 |
| *SbCMCDT | 5.0 | 2.5 | 5.0 |
| *JbCDTCM | 2.5 | 2.5 | 2.5 |
| *Jb~~CDT~~CM | 2.5 | 2.5 | – |
| *JbCDT~~CM~~ | 2.5 | 2.5 | – |
| *JbCM | 2.5 | n.p. | – |
| *DsCDTCM | 2.5 | 2.5 | 2.5 |
| *MpCDTCM | 2.5 | 2.5 | 2.5 |

The 24 reactions are listed in groups of four from A1-F4 with the corresponding substrate concentration. The enzyme concentrations used, denoted as […], are separately listed in the table on the right and are varied depending on the enzyme's individual activity and type of kinetic assay performed. The reaction time until quenching with either HCl or NaOH is shown in minutes. One hundred µL of 2 M HCl and 100 µL of 10 M NaOH were required for the CM discontinuous assay, whereas no HCl and 200 µL of 5 M NaOH were required for the CDT and the coupled CM+CDT discontinuous assays. Reactions that were not performed are indicated with 'n.p.' and assays that are not applicable with '–'.
